# Abundance redistribution increases predator-prey interaction potentials among North American birds

**DOI:** 10.64898/2025.12.19.695589

**Authors:** Heng-Xing Zou, Chia Hsieh, Phoebe L. Zarnetske, Kai Zhu, Brian C. Weeks

## Abstract

Species are shifting their geographical ranges, yet the consequences of these spatial redistributions for species interactions have not been examined at broad spatial and temporal scales. Here, we characterize shifts in the potential for predator-prey interactions among North American birds over the past 50 years. We first predict potential predator-prey interactions based on existing data, then construct a novel, abundance-weighted metric of interaction potential for each pair of predator and prey species in local communities. We found nearly universal increases in predator-prey interaction potentials across North America that are mostly driven by increases in predator abundances. The spatial redistributions of predators and prey lead to turnover of interactions within local communities that is less associated with changes in climate than with human-induced land use changes. Our results highlight the importance of considering traits, abundances, and species interactions—including known and potential interactions—when studying the ecological consequences of range shifts.

## Introduction

Across the world, species are shifting their geographical distributions in response to rapid environmental change (Fredston et al., 2025; Lenoir & Svenning, 2015; Parmesan & Yohe, 2003). Latitudinal and elevational shifts have been documented among plants, insects, amphibians, birds, and mammals under climate change (Chen et al., 2011; Lenoir et al., 2008; Moritz et al., 2008; Tingley et al., 2009); this spatial redistribution of biodiversity is likely to have far-reaching ecological consequences. While we have extensive evidence of the impacts of range shifts on community composition (La Sorte & Jetz, 2010; Lawlor et al., 2024; Zhu et al., 2024; Zou et al., 2025), our understanding of how range shifts have impacted species interactions remains largely constrained to isolated case studies (Grunsven et al., 2007; A. J. A. Stewart et al., 2015; J. E. Stewart et al., 2024). As range shifts alter community composition, and therefore the potential for species to interact with each other, applying a generalized framework to assess the effects of range shifts on this key ecological process is essential to reconstructing and predicting the effects of global change on species interactions and biodiversity.

When a species expands or contracts its range, its spatial overlap with other species changes, potentially resulting in changes to species interactions. In some cases, range expansion could result in novel interactions among species that have not encountered each other before, and range contraction could result in historical interactions disappearing (A. J. A. Stewart et al., 2015). Even when the bounds of a species’ spatial range do not change, variation in abundance within the range can affect the potential and strength of its interactions with other species (Chamberlain et al., 2014; HilleRisLambers et al., 2013; Vázquez et al., 2007). If one species becomes increasingly abundant, it is more likely to be encountered by the other species, increasing the potential for biotic interactions and vice versa. Among these changing interactions, predator-prey interactions are particularly important. For instance, the introduction of invasive alien predators can greatly impact native communities, driving native prey species locally extinct and affecting the fitness of native predators through competition (Dickman, 1996; Fagan et al., 2002; Hulme, 2017; Martin-Albarracin et al., 2015). These novel interactions can greatly change the energy flow of ecosystems, undermining biodiversity and the stability of food webs through changing top-down control (Lehman & Cáceres, 1993; Wainright et al., 2021).

Although climate change is typically considered a driver of range shifts (Chen et al., 2011; Lenoir et al., 2008; Moritz et al., 2008; Tingley et al., 2009), land use change can also contribute to shifts in spatial ranges, or the redistribution of abundances within species ranges (Lawlor et al., 2024). Land use changes can affect the connectivity, fragmentation, and overall quality of habitats, subsequently affecting both the distribution of species and the potential for certain interactions (Zarnetske et al., 2017). For instance, habitat degradation may lead to the contraction of species ranges, leading to less spatial overlap and therefore weaker interactions (Franco et al., 2006; Virkkala, 2016). On the other hand, more fragmented habitats may allow predators to have easier access to prey, strengthening the interactions (Andrén et al., 1985; Stirnemann et al., 2015). With the accelerating changes in climate and environment across the world, quantifying their combined effects on predator-prey interactions is important to understanding the ecological consequences of global change.

Despite their ecological importance, unlike abundance and community composition, species interactions are rarely examined at broader spatial and temporal scales (29). This may be because interaction data are notoriously difficult to collect: it is logistically impossible to observe every interaction happening in a natural system, and automated data collection remains challenging (Hortal et al., 2015; Strydom et al., 2021). For example, recent developments in joint species distribution modeling aim to capture the associations between species across space (32), but model outputs can only be interpreted as indicative of co-occurrence rather than biotic interactions between species (Blanchet et al., 2020; Zurell et al., 2018). Ideally, these statistical associations would be coupled with empirical data that clearly document interactions among species to enable understanding of the mechanistic role played by interactions in determining species distributions, but broad-scale validation is typically not possible due to limited field-collected interaction data. An alternative class of approaches is to use statistical models to predict possible interactions based on functional or phylogenetic information (Fricke et al., 2022; Llewelyn et al., 2023; Pichler et al., 2020; Strydom et al., 2021). For instance, ecological and life history traits such as habitat type, habitat density, and active times can determine the encounter probabilities for predators and prey (Hsieh et al., 2026), whereas morphological traits such as body mass - and the ratio of body masses of two species - and beak shape can determine the probability of successful predation (Fargallo et al., 2020).

We develop a novel approach to analyze the spatiotemporal shifts in known and predicted predator-prey interactions among North American birds, and to partition the importance of changing distributions of predators and prey in driving these shifts. The ecological importance of birds, along with the availability of large-scale datasets on bird functional traits (Tobias et al., 2022), abundances (Sauer et al., 2017), and a relatively comprehensive collection of predator-prey interactions (Hurlbert et al., 2021) makes them a powerful system for studying global change. The distributions and abundances of North American birds have changed significantly through time (Dupont & Dobson, 2025; Neate-Clegg et al., 2024), suggesting it is likely that over the past decades, there may have been important but undocumented changes in bird interactions. We examine changes in the potential for predator-prey interactions over the past half-century using abundance estimates that we generated across North America using the North American Breeding Bird Survey (NABBS) (Sauer et al., 2017). Leveraging known predator-prey interactions (Hurlbert et al., 2021), we developed a comprehensive set of potential predator-prey interactions based on key functional traits (Tobias et al., 2022; Wilman et al., 2014) by calculating the “functional compatibility” of potential prey species with different predators. We then define a probabilistic metric of “interaction potential” based on the functional compatibility and abundances of the predator and prey species, including recorded prey species and species predicted to be prey based on functional compatibility. Using this novel approach, we ask the following questions: (1) How have shifts in abundances affected potential predator-prey interactions over time? (2) What are the relative contributions of changing predator and prey relative abundances to changes in interaction potentials? (3) How do spatial distributions of predator and prey relative abundances and changes in predator-prey interactions connect to environmental variables?

## Methods

### Bird abundances and interaction data

We estimated changes in bird abundance using data from the North American Breeding Bird Survey (NABBS). NABBS is a long-term annual survey of birds in the United States and Canada via point counts at fixed sampling routes during the breeding season (May, June, or July; (Sauer et al., 2017)). We used NABBS data to estimate the abundance of species at a spatial scale of one degree latitude by longitude (hereafter a “local community”). This scale is based on the design of NABBS; it is the smallest spatial resolution that includes enough survey data to estimate bird abundances across the survey area (Zou et al., 2025). To generate our abundance estimates we used the 2022 release of the NABBS data and applied the following filters for data quality: (1) we removed years and regions that are data-deficient, resulting in a dataset spanning 1970-2021 (50 years of data; no survey conducted in 2020), (2) we only kept local communities with more than 10 species and that were north of 25°N, south of 55°N, and east of 135°W in the contiguous United States and Canada, and (3) we removed all species with insufficient data (< 50 records in total or recorded in fewer than three routes per local community in 1970-2021) or that are pelagic or nocturnal based on the EltonTraits 1.0 dataset (Wilman et al., 2014).

Next, we modeled the yearly abundances of all species in the filtered interaction dataset in each of the local communities by fitting hierarchical Bayesian models with a spatial intrinsic conditional autoregressive structure to raw count data with the R package *bbsBayes2* (Edwards et al., 2023). Model convergence was assessed using the Gelman-Rubin R-hat diagnostic with an R-hat threshold of < 1.1 (Brooks & Gelman, 1998). We corrected for the different detectability of species using established adjustment coefficients widely implemented for analyzing NABBS data (Dupont & Dobson, 2025; Rosenberg et al., 2019; Stanton et al., 2019; Zou et al., 2025). The relative abundance of each species in each year and local community was then calculated as the abundance of the focal species divided by the sum of abundances of the entire community (i.e., all species in the list of 470 species that are present in the local community in each year). To avoid confusion, we define “absolute abundance” as the raw abundance metric fitted from the model, as opposed to “relative abundance” calculated as above. See (Zou et al., 2025) for more information on the species selection, spatial filtering, model fitting process, model diagnostics, and detectability correction.

We obtained data on predator-prey interactions among birds from the Avian Diet Database, which compiled diet data from field observations or stomach content analysis of 759 primarily North American bird species (Hurlbert et al., 2021). We filtered the records to only those in which both the predator and the prey are birds and to only include interactions for species that are both within the list of 470 species from the BBS that met all our criteria for estimating relative abundance, resulting in a total of 14 predators and 251 prey species with 1561 interactions in 1196 local communities. Note that a bird species could be the predator in some interactions and prey in others. Since the location of predator-prey relationships was not included in the Avian Diet Database, we assumed that predator-prey interactions were the same across the species’ ranges. All scientific names were unified to the 2023 version of the Clements taxonomy (McTavish et al., 2025).

### Predicting predator-prey interactions

To identify likely predator-prey interactions not recorded in the dataset, we predicted potential interactions based on the “functional compatibility” between predators and potential prey. For each predator-prey pair, we calculated the similarity between the recorded prey of a predator and all other prey species in the dataset using cosine similarity (Hsieh et al., 2026; Xia et al., 2015) to identify species that are most functionally similar to the recorded prey. The higher the functional similarity to the recorded prey, the more likely that these species can be the potential prey of the predator. The cosine similarity of two prey species was calculated as the angular distance between their vectors in the multidimensional functional space given by their shared predators (Hsieh et al., 2026). Because a multidimensional functional space is constructed for each predator, the cosine similarity of the same pair of recorded and potential prey species can differ among predators. This improves the accuracy of predictions because the predator’s ecological information is also considered. Cosine similarity ranges from −1 to 1, with values closer to - 1 being less similar and values closer to 1 being more similar; a value of 1 would indicate identical traits.

To calculate cosine similarity, we compiled data on functional traits that are ecologically relevant to predator-prey interactions, including ecological, morphological, and nest traits. Ecological traits of prey, such as habitat, lifestyle, diet, and foraging, are descriptive of the activity patterns of species and are directly related to the potential of encountering predators (Fricke et al., 2022; Hsieh et al., 2026). Morphological traits such as body and appendage sizes directly affect whether a prey can be hunted and handled by potential predators and whether the predation will lead to a net gain of energy (Carbone et al., 2007; Fargallo et al., 2020). We obtained morphology, habitat, lifestyle, diet, and foraging traits from AVONET (Tobias et al., 2022) and EltonTraits 1.0 (Wilman et al., 2014). Although not specified, the Avian Diet Database may include incidences of nest predation, and we accounted for these common predation events by including traits on nest sites and structures (DeGregorio et al., 2016; Ibáñez-Álamo et al., 2015; Lima, 2009). Therefore, we included traits that describe nest structure, opening, and attachment (Chia et al., 2023). For each predator-prey pair, we calculated cosine similarity between the recorded prey and all 470 species in our species list. After resolving name changes, species mergers, and splits, the only truly missing species in many datasets was the Mexican Duck (*Anas diazi*). For the completeness of data, we imputed the functional traits for the Mexican Duck from the closely related Mottled Duck (*Anas fulvigula*), but the Mexican Duck was not in the Avian Diet Database. We did not impose additional filters on species ranges at this stage because the calculation of interaction potential filters out any species pairs without spatial overlap (see *Calculating and partitioning interaction potentials*).

There are two key challenges to this approach that require special consideration. First, we could not directly evaluate the accuracy using false positives (i.e., predicted interactions that have not been recorded) because these were very likely ecologically possible but unrecorded interactions due to limitations in the field-based data. Second, we could not split the data into a “training set” and a “testing set,” to then use the training set to predict interactions, and then match them to the testing set, because the functional space built from a subset of prey species for each predator would naturally be smaller and lead to poorer predictions. To overcome these challenges, we imposed several filters on the predicted prey species to improve the accuracy of predictions.

First, we only considered predicted prey with a cosine similarity larger than 0.95, which indicates very high functional similarity with the recorded prey species. If there are many predicted prey species for a recorded prey, we only considered the top 10 with the highest cosine similarity values. Second, we imposed a strict filter on habitat density (from open habitats such as desert, low shrubs, or the top of forest canopies to dense habitats such as lower-story forests or dense thickets) (Tobias et al., 2016, 2022): only potential prey with the same preferred habitat density as the recorded prey are considered functionally compatible with the predator. Third, we filtered the predicted prey species by the ratio of their body mass to the body mass of the predator species. While body mass ratio is important in determining compatibility between mammalian predators and prey (Carbone et al., 2007; Hsieh et al., 2026), in our dataset of observed avian predator-prey species pairs, we observed a wide range of body masses among the recorded prey of single predator species, suggesting body mass is likely not as strong a predictor of prey suitability in birds as it is in mammals (Figure S3). We therefore imposed a less strict filter on body mass based on the log mass ratio between potential prey and recorded prey:

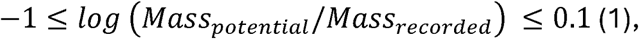

meaning that only potential prey with a body mass of 36.79% to 110.5% of the recorded prey are considered functionally compatible with the predator.

Our analyses assumed that functional traits and behaviors within species do not change over space and time. We tested the sensitivity of our results to this assumption using a sensitivity analysis that simulated spatial variation in IIP by assuming different thresholds of functional similarity among observed and predicted prey species in species-rich vs. species-poor communities. Specifically, predators in communities with higher prey species richness may be able to specialize on certain prey species with higher functional compatibility, while predators in species-poor communities may be forced to prey on species that are relatively less functionally compatible due to the lack of resources (Ferretti et al., 2020; Keppeler et al., 2025; Milana et al., 2016). Based on this simple assumption, we scaled the default threshold of functional compatibility (0.95) by species richness in community *i*(*N_i_*) across a range of values for a scaling factor *k*:

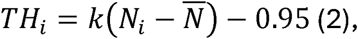

where *TH_i_* is the new threshold for community *i*, and *N̅* = 93.8 (range: 18-186) is the average species richness across all 1196 communities. We used an arbitrary *k* = 0.00025 to ensure that *TH_i_* would not be too close to 1 (which would filter out most predicted interactions) for the most diverse communities while not dropping too low for the least diverse communities. With this *k* value, the *TH_i_* ranges from 0.931 (least diverse community) to 0.973 (most diverse community). We did not find any qualitative differences in our main conclusions (Figure S7-S11, Table S4-S5).

### Calculating and partitioning interaction potentials

We assume that for predator-prey interactions to occur, the two species need to overlap spatially (i.e., have non-zero relative abundances in the same local community) and be functionally compatible. We defined the interaction potential between predator species 1 and prey species 2 at the grid *s* and year *t* as:

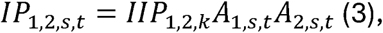

where *A_i,s,t_* is the relative abundance of species, and *IIP*_1,2_ is the intrinsic interaction potential between predator species 1 and prey species 2, calculated as the cosine similarity between the prey species 2 and a recorded prey (species *k*) of the predator species 1. If the interaction between predator and prey is recorded, we set *IIP* = 1, meaning that the two species are fully functionally compatible, and their interaction potential is solely determined by their probability of encounter within a grid *s* in a year *t*. If prey species 2 is functionally similar to *n* the recorded prey of predator species 1, then *IIP*_1,2_ is calculated as the average among all cosine similarities, i.e., 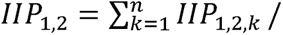 *n*. This metric provided a generalizable, broad-scale representation of predator-prey interactions that allowed us to characterize spatial and temporal shifts of the consequences of changes in predator and prey abundances.

Although relative abundances can serve as an approximation of encounter rates between the predator and prey, they may not accurately represent absolute abundances if the total abundance of the community also changes over time. Therefore, we used absolute abundances rather than relative abundances in Eqn. 3 to calculate interaction potentials, then conducted all downstream analyses. We found no qualitative changes in our results when using absolute abundances (Figure S12-S16 and Table S6-S7).

To better examine the contribution to temporal trends of interaction potentials between predator and prey, we further partition *IP*_1,2,*s,t*_ by log-transforming Eqn. 1:

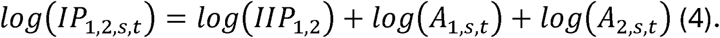

The temporal trend of the log-transformed interaction potential between predator species 1 and prey species 2 in the grid *s* can be modeled by a linear regression

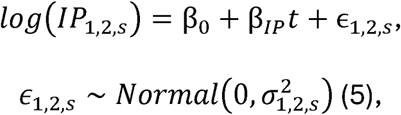

where *β*_1,2,*s*_ is the temporal slope, i.e., the temporal trend of the interaction potential. Writing *β*_1,2,*s*_ in the Ordinary Least Squares (OLS) solution, we have

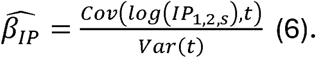

Because the intrinsic interaction potential does not change with time, Eqn. 5 can be written as

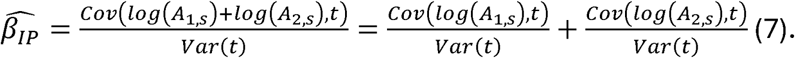

For species *i*, the temporal slope of its log-transformed relative abundance in the grid *s* can be similarly written in the OLS form,

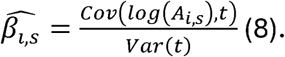

Combining Eqns. 6 and 7, the temporal slope of the log-transformed interaction potential between predator species 1 and prey species 2 is simply the sum of the temporal slopes of log-transformed relative abundances of species 1 and 2, i.e.,

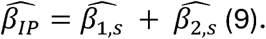

Although 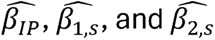 can be directly calculated by their respective OLS forms (e.g., Eqn. 7), we first fit linear models (Eqn. 5) to obtain information on statistical significance. We then selected species pairs with statistically significant temporal trends in interaction potentials after applying the Benjamini-Hochberg correction for multiple comparisons (Benjamini & Hochberg, 1995) and then decomposed these pairs by Eqn. 9. When fitting linear models, year was transformed by subtracting 1969 such that the first year in the data (1970) was 1.

### Overall predation risks for prey

One species may be the prey of multiple predators in each local community. To examine the temporal trends of the overall predation risk for each prey species, we calculated the predation risk as the sum of interaction potentials of all its predators. Then, we analyzed the predation risks of prey species by their functional groups (landbird, shorebird, waterbird, waterfowl, and other) and breeding biomes (arctic tundra, aridlands, boreal forest, coasts, eastern forest, forest generalist, grassland, habitat generalist, introduced, western forest, and wetland; see (Rosenberg et al., 2019) for complete definitions). For each grouping, we fit a linear mixed effects model:

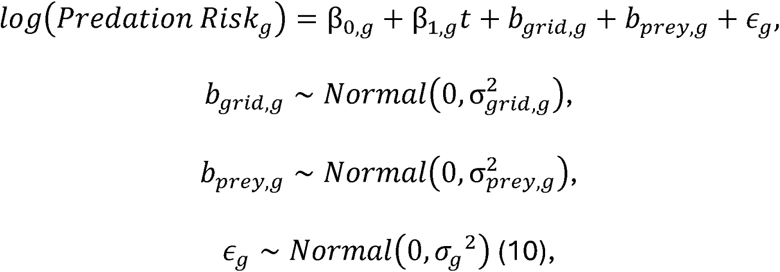

where *t* = *t_original_* – 1969 is the transformed year by subtracting the original years by 1969, such that the first year in the time series is 1. *b_grid,g_* and *b_prey,g_* represent the grid- and species-level random effects within each group of prey. We then corrected for multiple comparisons using the Benjamini-Hochberg method (Benjamini & Hochberg, 1995).

### Latitudinal shifts

To understand the spatial shifts in interaction potentials of each predator and prey pair, we quantified latitudinal changes in their abundances over time. For each species, the spatial and temporal smoothing of the model fitting process yielded continuous time series of abundance estimates for all its past and present spatial ranges from 1970 to 2021 (Edwards et al., 2023; Zou et al., 2025), meaning that the spatial range of each species does not change over time. In addition, species range limits in our dataset can be limited by both political borders and manual filters that are not ecologically meaningful. However, spatial redistribution of species abundance within local communities is a consequence of range shifts. For instance, if a species shifts northward, its abundance at the northern range edge will necessarily increase over time. To capture such shifts in latitudinal distributions of species abundance and interaction potential, we calculated the weighted mean latitude as the centroid of latitudinal distributions of each species’ relative abundance or each pair’s interaction potential. This measurement accounts for the distribution of abundances and can represent shifts in spatial ranges (Billman et al., n.d.; Brodie et al., 2025). For measurement *R*. (relative abundance or interaction potential) at a time *t*, the weighted mean latitude *y̅_t_* is calculated as:

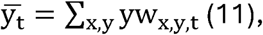

where *x* is the longitude and *y* is the latitude of each local community, and *w_x,y,t_* is the weight of each local community at latitude *y*, defined as:

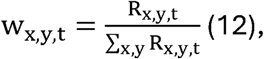

where the denominator is the sum of all measurements *R* across both latitudes *y* and longitudes *x* at time *t*.

We first calculated the latitudinal centroids of the entire ranges within our study area for each predator and prey species (i.e., range-wide latitudinal centroids). For any predator-prey interaction, the latitudinal centroid is defined by the spatial overlap of the predator and prey species, which may not be the entire spatial range of either species. We therefore also calculated the latitudinal centroids of the predator and prey’s spatial overlap, which are specific to each interaction pair. Finally, we calculated the latitudinal centroids of the interaction potentials for that pair. We estimated all temporal shifts in latitudinal centroids using a linear model with time:

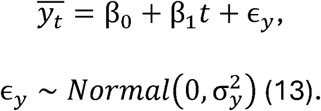

All p-values of linear models of predator and prey relative abundances and interaction potentials were adjusted by the Benjamini-Hochberg correction (Benjamini & Hochberg, 1995).

### Turnover of interactions and its association with environmental variables

We calculated the turnover of predator-prey interactions by considering the changes in their interaction potential between the beginning and the end of the time series (1970 to 2021). Because the model fitting process considers all past and present spatial ranges of each species over time (see *Latitudinal shifts*), the interactions in 1970 and 2021 for each local community consist of the same predator and prey species, but their interaction potentials are different due to changes in their relative abundances (i.e., changes in the weight of links between species). We first formatted these interactions as adjacency matrices with rows representing predators and columns representing prey species. We then calculated the turnover as the dissimilarity between the two interaction matrices in 1970 and 2021 using the Frobenius norm (‖A‖). This metric summarizes the difference between two interaction matrices in a holistic, single value that captures changes in all pairwise interaction potentials. Species with higher abundances will necessarily have higher interaction potentials, and the overall turnover value may be biased by changes in these abundant species. Therefore, we normalized the Frobenius norm using the interaction matrix at the beginning of the time series (1970). For each local community,

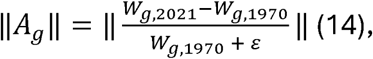

where *W_g_*_,1970_ is the interaction matrix for local community *g* in year 1970, and *ε* = 1^-10^ is a small offset to ensure a positive denominator. The numerator of the right-hand side of Eqn. 14 can be expanded as

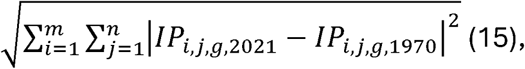

where *i* and *j* represent a pair of predator and prey, *m* and *n* represent the total number of predator and prey in the local community *g* (i.e., the dimensions of the interaction matrix), and *IP_i,j,s_*_,2021_ represents the interaction potential between predator *i* and *j* prey = in year 2021 and the local community *s*, calculated according to Eqn. 3. The turnover ‖A_*g*_‖ ranges from 0 to 1, with 0 meaning the communities have remained identical.

We then obtained historical estimates of land use data from the History Database of the Global Environment (HYDE) 3.3 dataset released in 2023 (Klein Goldewijk et al., 2017) for the years 1970 and 2021. This dataset includes anthropogenic land use types (“anthromes”) at a yearly resolution for years after 1950 generated from historical surveys, population censuses, and remote sensing data (Klein Goldewijk et al., 2017). We categorized land use types as the following: settlements, including dense settlements and villages; agriculture, including both croplands and rangelands; cultured, including residential and inhabited woodlands and drylands; wild, including wild woodlands and drylands (Ellis et al., 2010; Klein Goldewijk et al., 2017). These land use types represent areas of high- (settlements and agriculture), mid- (cultured), and low- (wild) levels of anthropogenic impact. We first calculated the proportion of each category in each local community (one-degree latitude and longitude grid cell) by aggregating the original data, which has a spatial resolution of 5 arc minutes, in the years 1970 and 2021, then subtracted the proportion of 1970 from 2021 to obtain the proportion of land use changes. For some local communities and land use types, these differences are 0, meaning no change occurred between 1970 and 2021. For each local community, the differences of these land use types add up to 0 and are therefore tightly correlated. To avoid this collinearity, we dropped the land use type that contributed the least to the turnover values, whose effect will be combined with the intercept of the linear model. All fitted slopes should then be interpreted as effect sizes of each environmental factor relative to the dropped variable. To assist with the interpretation, we visually examined the linear relationship between the turnover values and each of the four land use types, then removed agriculture, the one with the flattest slope (i.e., closest to 0, indicating the smallest effect; Figure S4). To compare the relative importance of land use changes and climate change, we included differences of mean annual temperature and precipitation between 1970 and 2021 in each local community (i.e., a single difference value for each 1-degree grid), and their interaction terms. These bioclimatic data were obtained from WorldClim2 (Fick & Hijmans, 2017) and aggregated to each local community (one-degree latitude and longitude grids).

Finally, the turnover of interaction potentials was caused by changes in species relative abundances. When fitting relative abundances, we used a hierarchical Bayesian model with spatial information (see *Relative abundances and interaction data*), which would leave a spatial signature on the abundances, with local communities close to each other having similar abundances. We corrected for this spatial autocorrelation by fitting a linear model with spatial lag using the R package *spdep* (Bivand, 2022). Specifically, the turnover (dependent variable) of the focal community could be influenced by its neighbors; this was close to the spatial process incorporated when fitting species abundances. Although the model fitting process considered only directly adjacent neighbors (Edwards et al., 2023), based on the residual analysis of a non-spatial linear model (E. J. Pebesma, 2004), the extent of spatial autocorrelation was much larger than this distance (Figure S6). To balance this large extent while mostly capturing the spatial process of the abundance models, we chose a slightly larger extent of all neighbors within a five-by-five grid (i.e., 24 neighbors) centered at each local community. Writing out the full model:

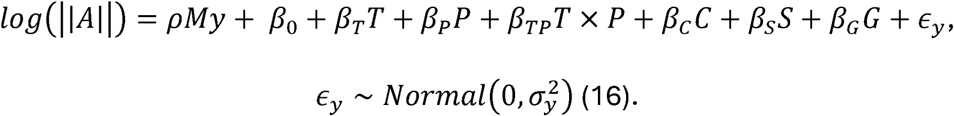

The response variable of a single local community, turnover ||*A*||, is log-transformed to account for the skewness in its distribution. *ρMy* is the spatial dependence term, where *ρ* is the spatial lag coefficient, *M* is the spatial weights matrix that defines the closest 24 local communities from the focal community, and *y* is the average turnover value of all the neighbors. We chose a relatively large number of neighboring communities because *T* and *P* are changes in mean annual temperature and precipitation, and *T*×*P* represents their interaction. *C*, *S*, and *G* represent changes in proportions of cultured, settlements, and agricultural lands. Because agriculture is removed to avoid collinearity *β_C_*, *β_S_*, *β_G_*, and their corresponding statistics are interpreted as the difference from the mean effect of changes in agricultural land. Because this effect of agricultural land use change was close to 0, *β_C_*, *β_S_* and *β_G_* can be interpreted close to their numeric values. All environmental variables were scaled to the mean in model fitting.

We conducted all analyses in R version 4.4.1 (R Core Team, 2024) with packages *tidyverse* (Wickham et al., 2019) and *sf* (E. Pebesma, 2018).

## Results

### General increases in predation potential driven by predators

Across North America and nearly all predator species, we found general increases in the modeled potential for predator-prey interactions over time (Figure 1-2). There were spatially clustered hotspots of change, with the greatest number of pairs with changing predicted interaction potentials (positive or negative) through time concentrated in the Northern Rockies, Great Basin, Prairie, and Appalachian Mountains (Figure 1A; Figure S1). These patterns are generally consistent when considering only recorded interactions or including both recorded and predicted interaction pairs (Figure S1). These temporal and spatial patterns reflect substantial change in species relative abundances within communities.

**Figure 1.**
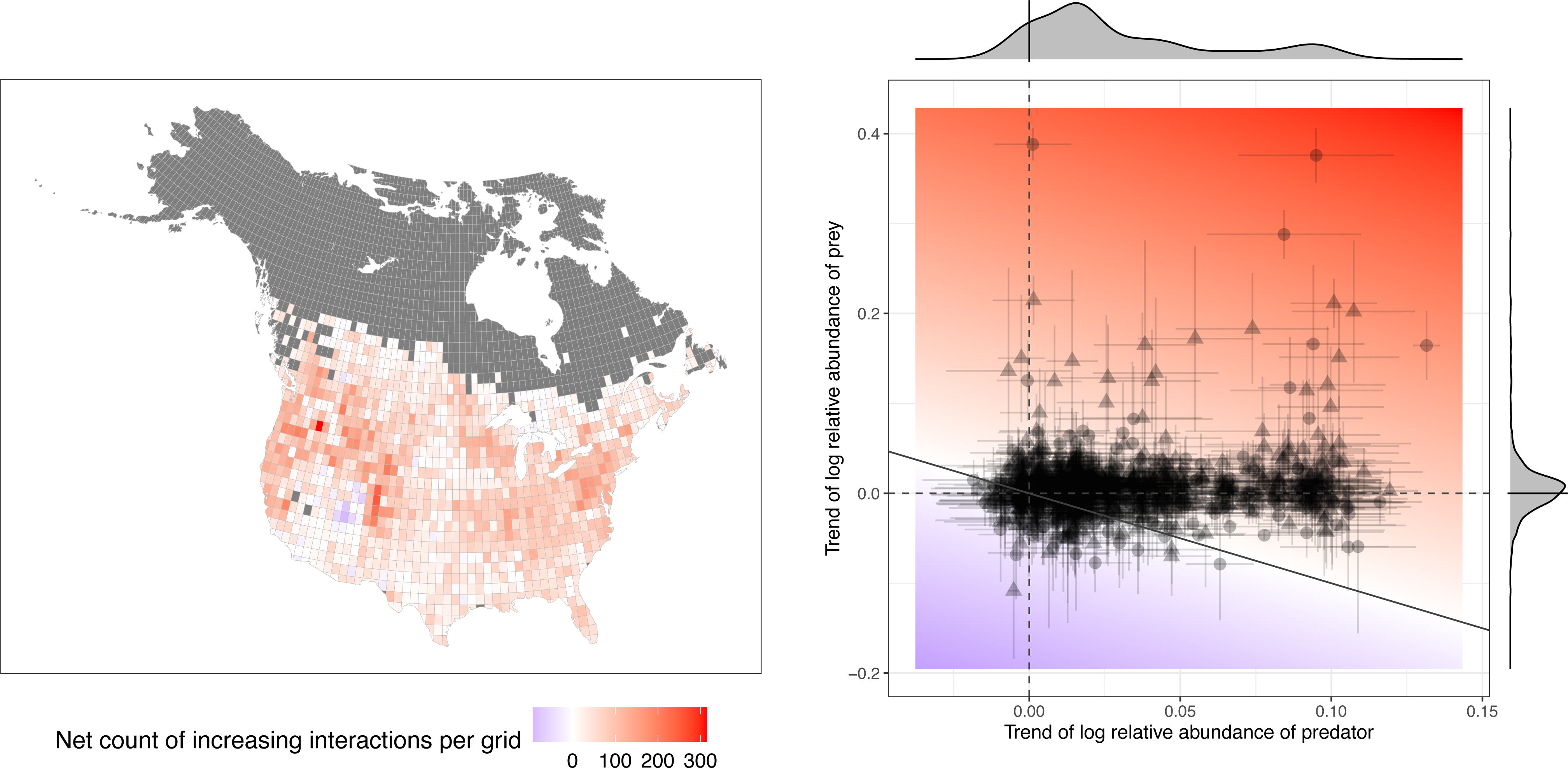
Most pairwise predator-prey interactions have increased their potential over time due to the increasing relative abundance of predators, not prey. Panel A (left): map showing the net count of increased interaction potentials, calculated by the number of pairs with increasing interaction potentials minus that of decreasing potentials in each one-degree latitude and longitude grid. Panel B (right): contribution of predators and prey to temporal changes in their interaction potentials, for all predator-prey pairs and in all local communities. For each pairwise interaction, the x-axis represents the mean temporal trend of predator relative abundances, and the y-axis represents the mean temporal trend of prey relative abundances, with associated standard deviations. Pairs above the 1:1 line indicate increasing interaction potentials over time (i.e., positive temporal trends). Marginal density plots represent the distribution of temporal trends. Both panels included interactions recorded in the existing dataset (triangles) and interactions predicted by the “functional compatibility” (dots). For both panels, one species pair will be counted multiple times if its interaction potential significantly increases/decreases in multiple local communities.

**Figure 2.**
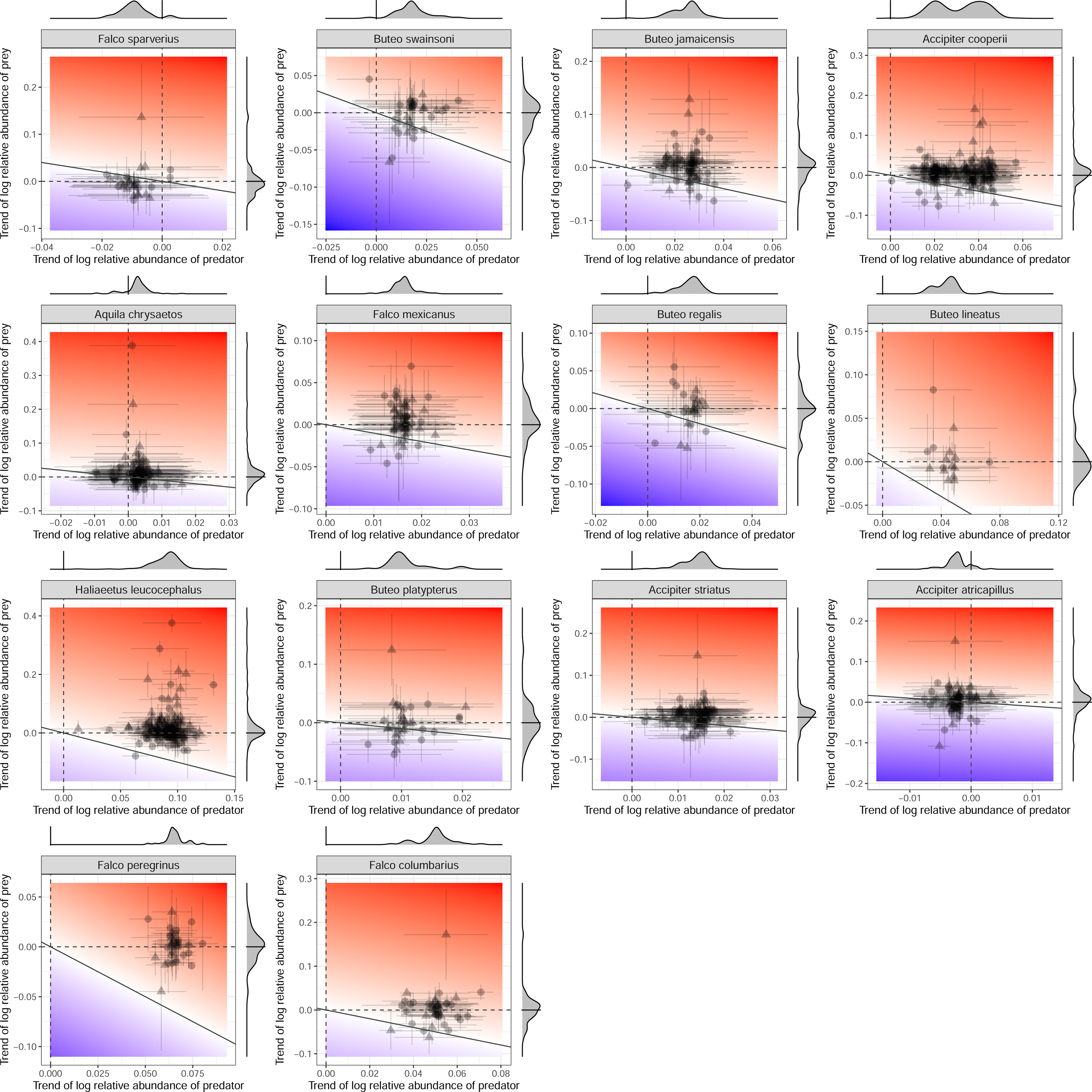
Contribution of predators and prey to temporal changes in their interaction potentials, for all pairs across all local communities by predator species. For each pairwise interaction, the x-axis represents the mean temporal trend of predator relative abundances, and the y-axis represents the mean temporal trend of prey relative abundances, with associated standard deviations. Pairs above the 1:1 line indicate increasing interaction potentials over time (i.e., positive temporal trends). Marginal density plots represent the distribution of temporal trends. All panels included interactions recorded in the existing dataset (triangles) and interactions predicted by the “functional compatibility” (dots). One species pair will be counted multiple times if its interaction potential significantly increases/decreases in multiple local communities.

By partitioning the temporal trends of each pairwise predator-prey interaction into changes in predator and prey abundances over time, we found that the widespread increase in predator-prey interaction potentials in local communities was driven predominantly by increases in predator relative abundances in local communities. The temporal trends of prey relative abundances center close to zero, meaning that, in general, prey abundances are relatively stable over time (Figure 1B). These trends are shared among all predators except American Kestrel (*Falco sparverius*) and American Goshawk (*Accipiter atricapillus*), which have declined in relative abundances across almost their entire ranges (Figure 2, density plots on the margin of the x-axis; see also Table S6). These results are consistent whether only recorded interactions are used or both recorded and predicted interactions are used (Figure 1, Figure S2).

To understand how shifts in potential predator-prey interactions might impact prey species, we examined the combined predation risk faced by each prey species by summing all their interaction potentials within each year and local community, then modeled the relationship between prey predation risk and time grouped by their breeding biomes and functional groups. We found that all groups of prey species show increasing predation risk over time, but waterfowl (ducks, geese) and waterbirds (shorebirds, wading birds) show the largest increases (Figure 3A; Table S1). Relatedly, prey breeding in coastal and wetland habitats, which are mostly waterfowl and waterbirds, have experienced the largest increase in predation risk (Figure 3B). Together, our analyses reflect an increasing potential for predator-prey interactions in North America through time, mostly driven by the increasing abundance of predators.

**Figure 3.**
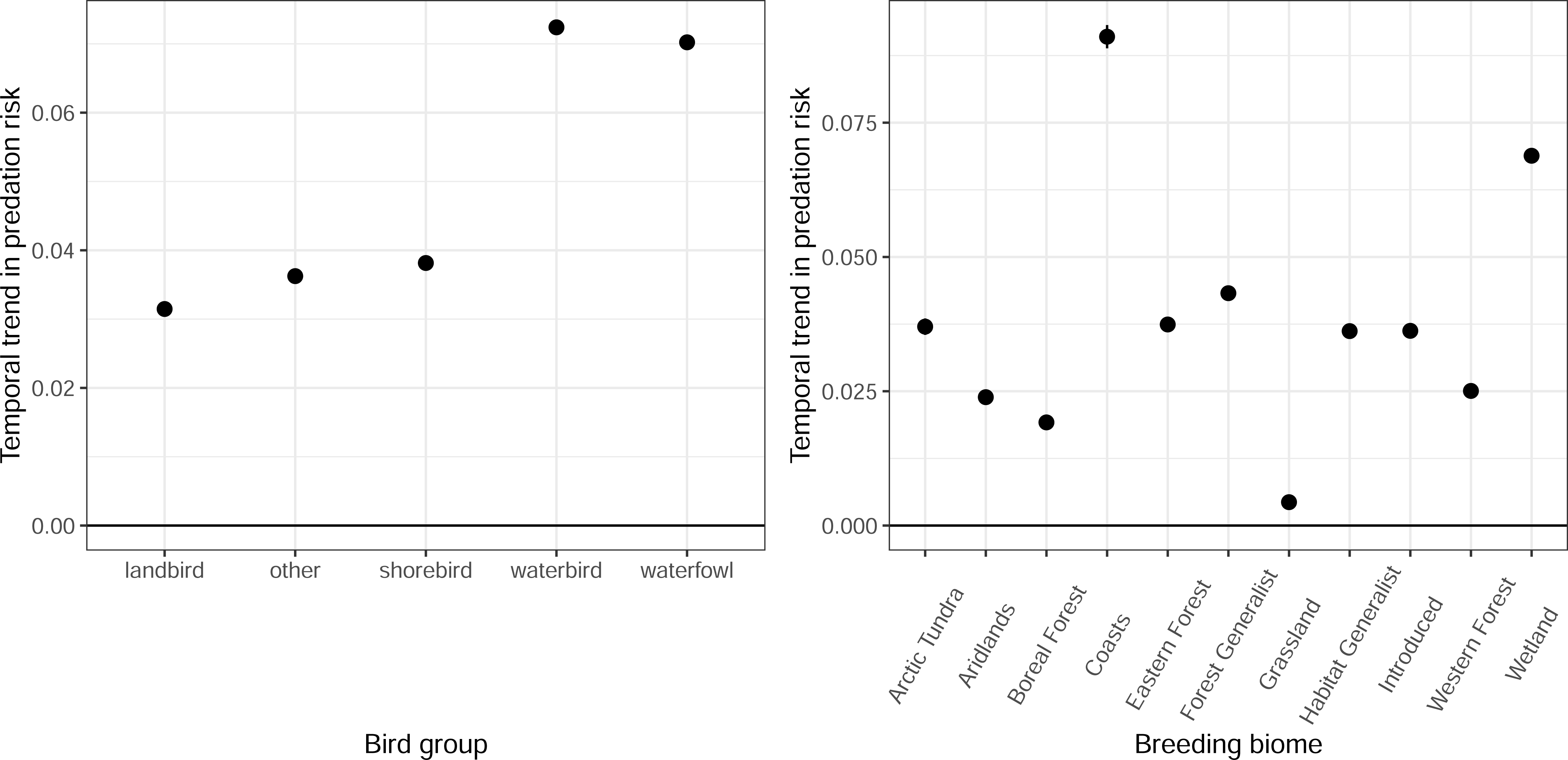
Temporal trends of overall predation risks by functional groups and breeding biomes of recorded and predicted prey species; points and lines indicate effect sizes and standard errors. Panel A (left): temporal trends by functional group; panel B (right): temporal trends by breeding biomes. Non-significant effects are shown in lighter shades.

### Latitudinal shifts among predators and prey

Out of 14 predator species with sufficient data (predators that have more than 10 recorded prey species), eight (57.1%) showed significant northward shifts of latitudinal centroids of their entire range, five (35.7%) showed significant southward shifts, and 1 showed no significant shift (Figure 4; Table S2). Out of the 370 prey species, 180 (48.6%) showed significant northward shifts, while 122 (33.0%) showed significant southward shifts, and 68 showed no significant shifts. Out of 1197 documented and predicted predator-prey interactions, 532 (44.4%) showed significant northward shifts, while 413 (34.3%) showed significant southward shifts; these proportions are not significantly different among recorded and predicted interactions (chi-square test, df = 1, *p* = 0.125). Latitudinal shifts in abundance altered the latitude of the abundance-weighted range centroid for the predator (Figure 4). These changes in abundance can be accompanied by a correlated latitudinal shift in interaction potentials between the predators and their prey. For instance, as a predator’s abundance shifted northward, its interaction potential could also shift northward. However, this was not necessarily the case since we observed considerable heterogeneities in these latitudinal shifts. For instance, the southward shift in the relative abundance centroid of the Bald Eagle (*Haliaeetus leucocephalus*) was much stronger than the shifts in most of the potential predator-prey interactions in which it participates. Similar mismatches may indicate a weak alignment of latitudinal shifts in predators and their avian prey. Overall, the latitudinal centroids of predator relative abundance, prey relative abundance, and potential of predator-prey interactions have shifted more towards the north, generally corresponding to the expectations under a warming climate, but with large variation among species.

**Figure 4.**
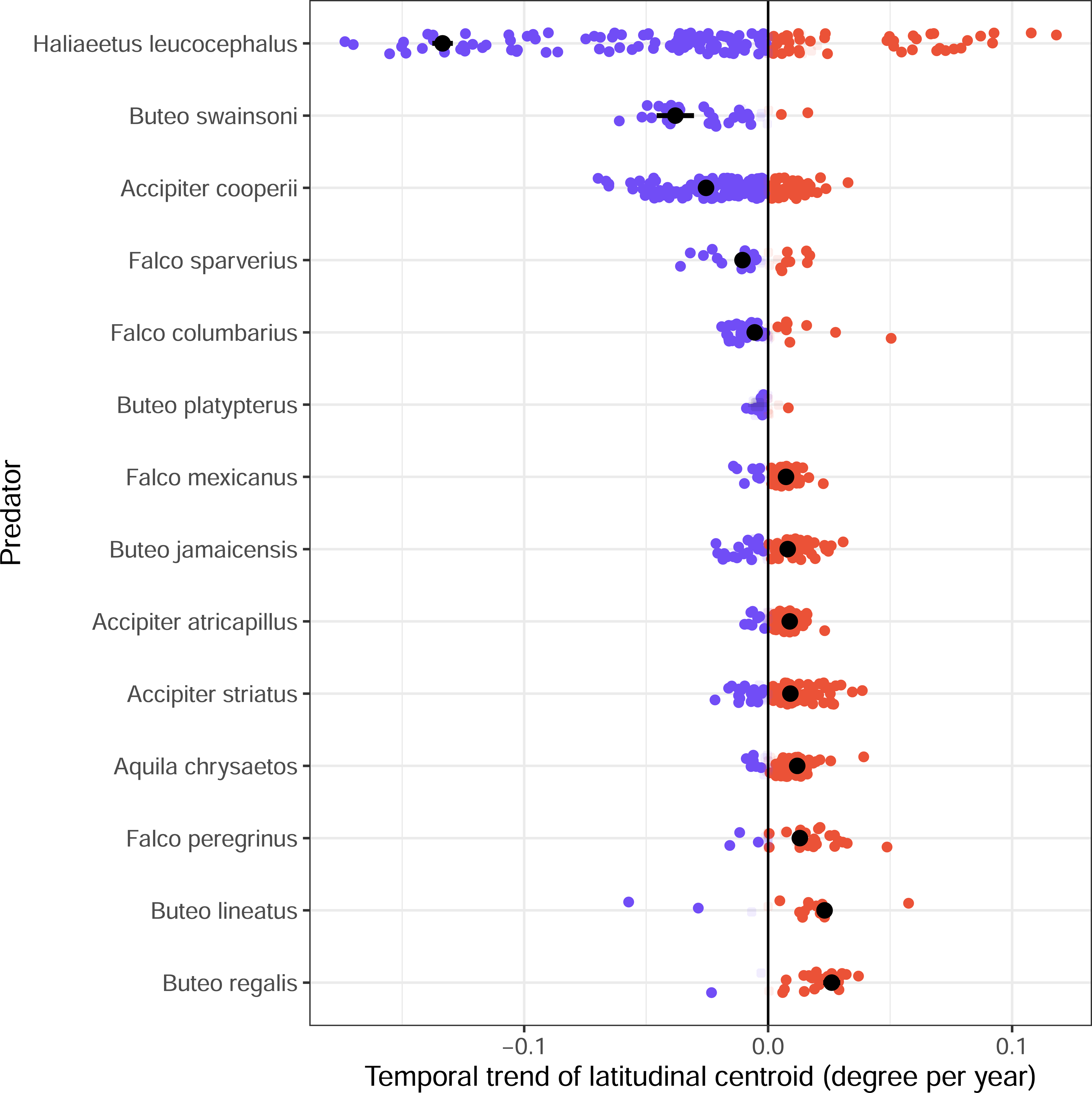
Latitudinal shifts over time of predator relative abundances in their entire ranges (black dots), and in their spatial overlap with their prey species (colored dots), including both recorded and predicted interactions. For a predator, because the spatial overlap with each prey species is different, each predator-prey pair will have an individual latitudinal shift. On the x-axis, positive numbers indicate northward shifts, negative numbers indicate southward shifts. Non-significant shifts are shown in lighter shades.

Next, we explored the contributions of shifting distributions of the predators and the prey to the latitudinal shifts in their interaction potentials. Visually examining the latitudinal shifts for each predator, we found that they follow two patterns (Figure 5). First, the abundance centroid of the predator shifts north or south, while the overall abundance centroids of all its prey species do not show any strong trends, including Bald Eagle, Cooper’s Hawk (*Accipiter cooperii*), Red-shouldered Hawk (*Buteo lineatus*), Swainson’s Hawk (*Buteo swainsoni*), American Kestrel (*Falco sparverius*), and Merlin (*Falco columbarius*). For these predators, the spatial distributions of their interaction potentials do not show visible latitudinal shifts towards either direction. Because the predators have shifted significantly towards the south, this lack of a clear visual pattern indicates the latitudinal shifts of predator interaction potentials may be more dependent on the latitudinal shifts of prey rather than predators. Second, both the predator and its prey species show trends of shifting northward, including Golden Eagle (*Aquila chrysaetos*), Sharp-shinned Hawk (*Accipiter striatus*), American Goshawk (*Accipiter atricapillus*), Red-tailed Hawk (*Buteo jamaicensis*), Ferruginous Hawk (*Buteo regalis*), Prairie Falcon (*Falco mexicanus*), and Peregrine Falcon (*Falco peregrinus*). In this case, disentangling the contributions of the predator and prey species was difficult. In our dataset, we did not find scenarios in which both predator and prey shifted south. One predator, Broad-winged Hawk (*Buteo platypterus*), did not shift significantly in either direction, but it also had fewer interactions (Figure 5). In general, contrary to the working hypothesis of a northward latitudinal shift under a warming climate, predators have shifted their abundances both northward and southward.

**Figure 5.**
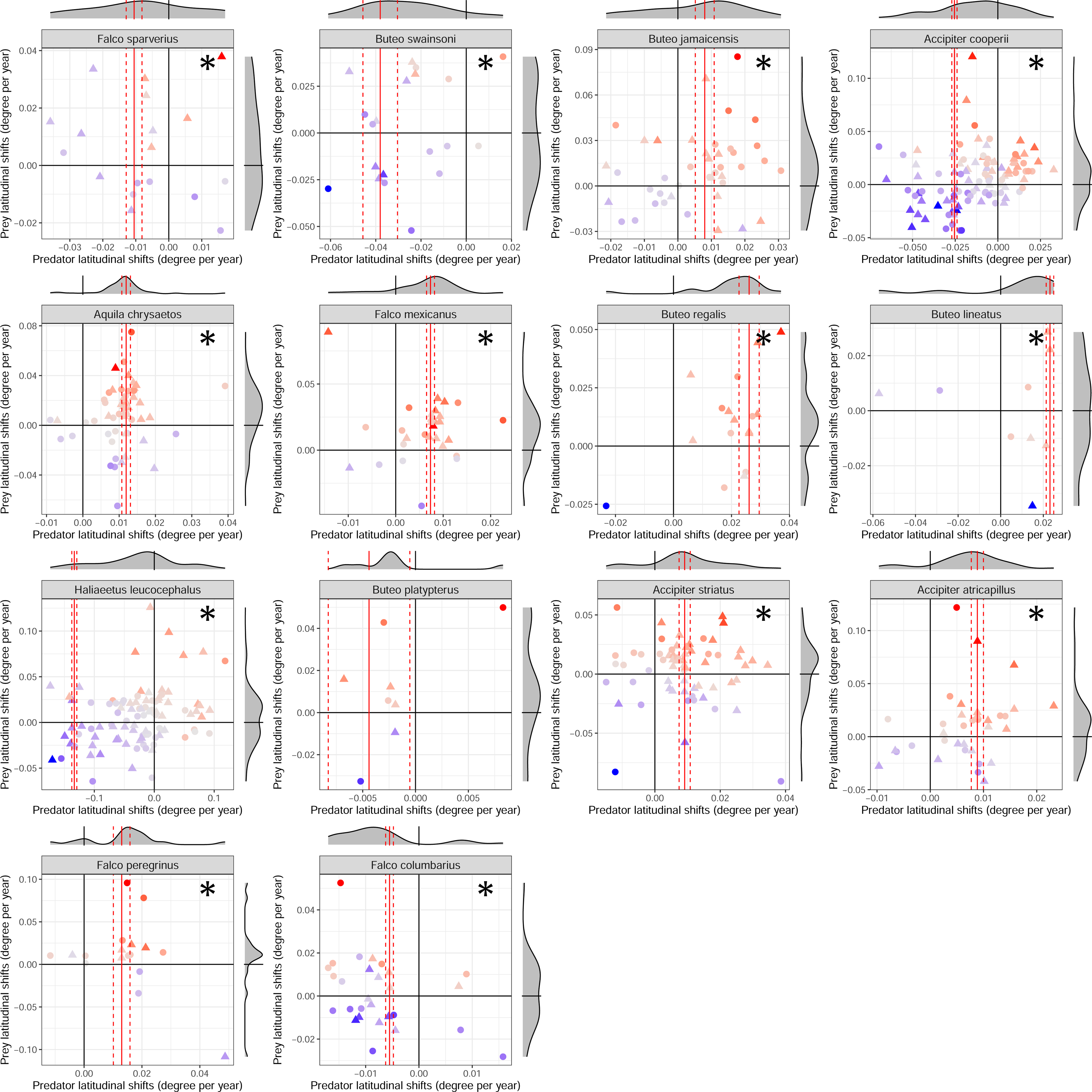
Latitudinal shifts of interaction potentials, decomposed by the latitudinal shifts of predators and prey relative abundances. Each panel represents both recorded and predicted interactions of one predator. The x- and y-axes indicate the latitudinal shifts of predators and prey, respectively, with positive values indicating northward shifts, and the marginal density plots show the distribution of latitudinal shifts, with black solid lines indicating 0 (no shift). Colors of each dot indicate the latitudinal shifts of each pairwise interaction potential (red: shifting significantly north; blue: shifting significantly south), dots indicate predicted interactions, and triangles indicate recorded interactions. The red vertical lines indicate the fitted slope and standard errors of the overall latitudinal shift of the predator, and the asterisks indicate that the means of the distributions of shifts in predator interaction potentials are significantly different from zero.

### The role of human-induced land use changes

Many local communities have zero or low changes in land use (Figure S4-S5), making the patterns mostly driven by a small number of communities with drastic changes. Visually inspecting the maps, settlements and cultured lands have increased in the Midwest and the Southeast, reflecting urbanization and suburbanization, whereas agricultural land decreased and wild land increased along the Gulf Coast and in southern Florida (Figure S5). Despite this spatial variation, we found significant positive associations between turnover of predator-prey interaction potential and changes in settlements (urban and suburban developments; slope = 0.315, 95% confidence intervals = [0.162, 0.468], *p* < 0.0001; see Table 1 for all slopes and statistics), meaning that increasing urbanization and suburbanization are associated with changes in predator-prey interactions. On the other hand, neither change in mean annual temperature, precipitation, nor their interaction was significantly associated with the turnover. These results are qualitatively similar when we only consider observed predator-prey interactions (Table S3). Overall, the relationships between land use changes and the turnover of predator-prey interactions were stronger than the relationship between turnover and climate change. Because turnover was entirely due to changes in relative abundances in local communities, our results highlight the impact of land use changes on bird population changes within ranges.

**Table 1.** Statistics of the linear model with spatial lag between turnover of predator-prey interactions and environmental changes between the years 1970 and 2021, calculated from relative abundances. Turnover was calculated as the difference between matrices of interaction potentials, including both recorded and predicted predator-prey interactions. Environmental change includes changes in land use, mean annual temperature, and mean annual precipitation, and the interaction between changes in temperature and precipitation within each local community. See *Methods* for details on model fitting.

| Environmental change | Estimate | SE | z value | p value |
| --- | --- | --- | --- | --- |
| (Intercept) | 0.898 | 0.194 | 4.618 | <0.0001 |
| Temperature | 0.040 | 0.059 | 0.688 | 0.492 |
| Precipitation | 0.017 | 0.069 | 0.244 | 0.807 |
| Settlements | 0.315 | 0.078 | 4.026 | <0.0001 |
| Cultured | -0.045 | 0.126 | -0.361 | 0.718 |
| Agriculture | -0.075 | 0.127 | -0.593 | 0.553 |
| Temperature: precipitation | 0.053 | 0.061 | 0.869 | 0.385 |
| N = 1179, df = 1170; $\rho = 0.884$, p value for $\rho < 0.0001$; pseudo $R^2 = 0.463$ | | | | |

## Discussion

Drastic changes in both the abiotic and biotic environments are expected to result in extensive spatial redistribution of species abundances that could greatly affect the dynamics of biotic interactions. Using 50 years of bird abundance data in North America, we found a nearly universal increase in the potential for interaction between predator and prey species. Across all but two of the predator species, these trends are predominantly driven by the increasing abundances of the predators. The increased abundances of predator species can lead to stronger top-down control on the bird community and other trophic levels. These results highlight how the spatial redistribution of birds could affect not only the composition, but also species interactions within communities.

The nearly universal population increase of birds of prey in North America over the past half-century has been well documented by long-term surveys (Hoffman & Smith, 2003; Rosenberg et al., 2019). Most of these increases have been attributed to two anthropogenic drivers. First, conservation policies in the 1960s and 1970s have led to large increases in the populations of many birds of prey. For instance, the banning of DDT increased the reproductive success of many species, assisting the rebound of their populations (Padayachee et al., 2023; Ratcliffe, 1967). Some species, such as the Peregrine Falcon and Bald Eagle, have also received intense, targeted conservation efforts (Bowerman et al., 1995; Kauffman et al., 2003). Because many of these policies coincide with the start of the North American Breeding Bird Survey (NABBS), it is not surprising that we find increasing relative abundances of birds of prey in North America across the NABBS data. Second, many birds of prey have benefited greatly from human-induced land use changes. Species such as Cooper’s Hawk and Peregrine Falcon have increased in urban and suburban areas where they are well-suited to take advantage of resources like abundant prey, such as Rock Pigeon (*Columba livia*), and smaller passerine species at backyard feeders (Mannan & Steidl, 2018; Pagel et al., 2018; Rosenfield et al., 2018).

Despite the general increases in the potential interactions between predator and prey species, we found significant but often idiosyncratic latitudinal shifts of predators, prey, and their interactions. Latitudinal shifts of avian predators at a continental scale may reflect the importance of multiple factors, including climate, environment (e.g., land use change), conservation efforts, and other potential mechanisms such as the ecology of each species. For instance, climate-induced range shifts have been widely documented in birds (Neate-Clegg et al., 2024; Tingley et al., 2009), especially among passerine species, but we found mixed evidence of this in the predator species, with five of the 14 predators shifting southward instead of northward. In some cases, it does seem likely that climate change may be driving shifts in interaction potentials, e.g., patterns that are consistent with predators tracking range shifts of their prey (Lawlor et al., 2024): the distribution of Sharp-shinned Hawk has shifted significantly northward, and this may arise from the spatial tracking of its prey, which are mostly small passerine species that have undergone northward shifts (Neate-Clegg et al., 2024). However, when considering the turnover of predator-prey interactions within entire local communities, we found weak effects of changes in climate, with stronger signals of human-induced land use changes, especially increases in urban and suburban areas. Furthermore, our analyses did not explicitly evaluate whether latitudinal shifts in predators arise from the spatial tracking of prey. Although our analyses are consistent with the hypothesis that some predators may be tracking certain avian prey species, these relationships are suggestive and could be more fully explored in future studies that explicitly test for such tracking using abundance and environmental data on finer spatial and temporal scales.

The inconsistent latitudinal shifts may also reflect a greater significance of idiosyncratic drivers of abundance and distribution for each species, even among species that are known to share similar environmental drivers of their population dynamics. Cooper’s Hawk and Bald Eagle, two species that have greatly benefited from recent urbanization and targeted conservation efforts, have both shifted significantly southward, whereas another similarly recovering species, the Peregrine Falcon, has shifted northward. These differences may arise from ecological differences among the individual predator species that we did not explore in our analyses. This may be related to one important limitation in our study: we only considered the avian prey of all predators, while the predators differ in the proportion of non-avian prey in their diet. For example, the diet of the Bald Eagle contains 46.8% fish, whereas mammals constitute 29.6% of the diet of the Cooper’s Hawk (both by weight/volume in stomach content; see (Hurlbert et al., 2021)). Therefore, the latitudinal shifts of some predators might arise from the potential tracking of non-avian prey such as fish and small mammals, which can also be impacted by climate and land use change (Comte & Grenouillet, 2013; Fredston et al., 2025; Moritz et al., 2008), or even from a dynamic shift of the prey species (Bourbour et al., 2024; Tornberg et al., 2014) that may lead to changes in ratios of non-avian prey. Conversely, our calculations of the interaction potentials likely represent an upper bound of predation risk, as most of the predators analyzed regularly obtain resources from non-avian prey. Taken together, our results highlight the complexity of mechanisms of bird range shifts and the need for more in-depth analyses in identifying these mechanisms among predator species.

Despite the overall increase in interaction potentials across North America, there is significant spatial variation that reflects different processes across biogeographical regions. For instance, major mountain ranges such as the Rocky Mountains and Appalachian Mountains appeared to show a higher number of increasing interaction potentials. This is consistent with the hypothesis that elevational range shifts are more likely to increase species interactions because of the clustering of species within smaller areas towards mountain tops ((Carlson et al., 2022; Freeman et al., 2022; Lawlor et al., 2024); but see (Elsen & Tingley, 2015)). Our methods provided a general estimation of interaction potentials on broad spatial and temporal scales, enabling robust comparisons across North America. However, although we calculated interaction potential between predator and prey based on their functional compatibility and abundance, with the coarse 1-degree spatial resolution in our analyses, we could not explicitly incorporate other mechanisms governing predator-prey interactions such as the fine-grained environmental factors, such as microhabitats and spatial refuges, and things important to behavioral ecology such as perception, previous exposure, and adaptation (Hein & Martin, 2020; Martinez-Garcia et al., 2020; Schmitz, 2017; Sih et al., 2010). Future studies could analyze high-resolution abundance estimates or tracking data and environmental data to better understand potential effects of environmental changes on encounter rates over space and time (Furey et al., 2018; Mirski et al., 2024; Yanco et al., 2025) to investigate the spatial patterns observed in our study in detail.

Increasing interaction potentials have multiple possible drivers, each with different ecological implications. We attribute the increased interaction potentials largely to the increasing abundance of predator species. Birds of prey contribute to the regulation of the populations of smaller birds and rodents, and increased top-down control is expected to have multi-trophic effects, potentially cascading through the community (Terraube & Bretagnolle, 2018). The ecological consequences of such trophic cascades can be complex and depend on the specific species being affected. Some of these impacts are likely to affect important ecosystem services; for example, increasing predation on insectivorous birds may lead to an increase in leaf damage by insects (Mäntylä et al., 2011). Understanding these processes is thus of both ecological and conservation significance. North American avifauna have faced steep declines in abundance over the last 50 years, with all but a few major groups showing dramatic decreasing trends (Rosenberg et al., 2019). Our results confirm this general increase in the abundance of birds of prey. The increase in predator abundance coincides with the decrease of most of their avian prey, but there is no evidence indicating a causal relationship. Rather, the decline in major prey groups such as smaller passerines is likely due to a combination of climate change, habitat loss, resource depletion, and contamination (Dupont & Dobson, 2025; Probst et al., 2026; Rosenberg et al., 2019; Spiller & Dettmers, 2019). The ongoing decline of these prey groups may further affect the fitness of predators, especially those specializing in avian prey species. Further analyses are necessary to understand the ecological impact of increasing predators and declining prey species.

Studying the ecological consequences of species redistributions requires data on biotic interactions. In our analyses, we expanded previous field-collected data on predator-prey interactions by predicting the potential interactions among functionally compatible predator and prey species. Then, we developed an interaction potential index based on this functional compatibility and abundances, which is a rough estimate of encounter probability without involving predictions from climatic niche modeling. Our index of interaction potential provides a parsimonious, holistic, probabilistic measurement that can be applied to quantifying all types of interactions between spatially and temporally overlapping species. The performance of this metric depends on the quality of the input data, including the specificity of the interaction data and the corresponding selection of functional traits (Strydom et al., 2021). For instance, we selected nest traits of prey in predicting interactions because nest predations are common and associated with nest sites and structures (DeGregorio et al., 2016; Ibáñez-Álamo et al., 2015; Lima, 2009), but the predator-prey interaction dataset we used does not contain detailed information such as the prey’s life stage (Hurlbert et al., 2021); our analyses were also constrained by the lack of high-resolution spatial and temporal variations in species interactions and functional traits. Interaction data with broader coverage and higher resolution are necessary to further advance the prediction of species interactions and our understanding of how they change over space and time.

Using a unique analytical pipeline that captures both observed and predicted interactions, we find that there have been broad trends toward increased potential interactions among predator and prey species in North American birds. Our results reflect the importance of considering the impacts of spatial shifts in abundance on trophic dynamics across broad spatial and temporal scales. Moving forward, it will be critical to better understand the ecosystem-level consequences of these shifting dynamics.

## Acknowledgements

We thank members of the Weeks and Zhu labs and IGCB Postdoc Fellows for their feedback on the project. We thank the numerous volunteers who collected data for the North American Breeding Bird Survey. H.-X.Z was supported by the Postdoctoral Fellowship from the Institute for Global Change Biology at the University of Michigan. C.H. was supported by the Presidential Postdoctoral Fellowship at Michigan State University. K.Z. was supported by the National Science Foundation award 2306198 and USDA McIntire-Stennis Capacity Grant award 25-PAF01509. B.C.W. was supported by the David and Lucile Packard Foundation. This research was supported in part through computational resources and services provided by the Advanced Research Computing at the University of Michigan, Ann Arbor.

## Authors contributions

H.X.-Z. conceived the idea, curated and analyzed data with input from C.H., P.L.Z, K.Z., and B.C.W. C.H. contributed to the prediction of predator-prey interactions. H.X.-Z. wrote the first draft, and all authors contributed to subsequent review and editing.

## Data and materials availability

All primary data are publicly available, and all generated data will be permanently archived upon acceptance. Code is available on GitHub: https://github.com/hengxingzou/Zouetal2025b_bioRXiv and will be archived upon acceptance.

