## Supplementary Material for "Abundance redistribution increases predator-prey interaction potentials among North American birds"

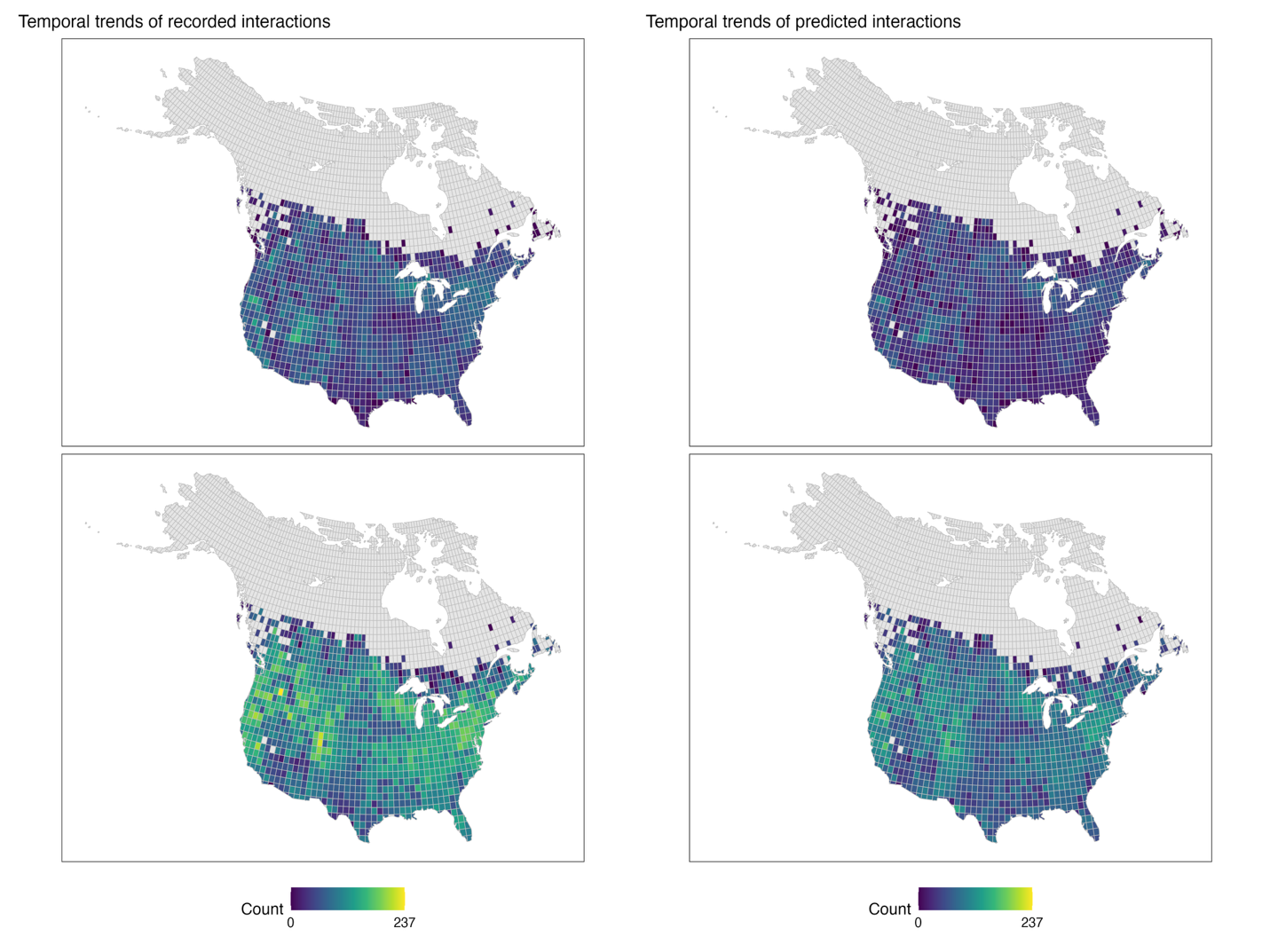

**Figure S1.** Map of temporal trends of interaction potentials in each local community. Top row: number of predator-prey interactions with decreasing interaction potential; bottom row: number of predator-prey interactions with increasing interaction potential. Left column: interactions recorded in the existing dataset; right column: interactions predicted from the existing dataset. Only species pairs with statistically significant temporal trends of interaction potentials are included. Numbers are sums of species pairs across all local communities: if a species pair shows significantly increasing interaction potential in *n* local communities, it will be counted *n* times.

**Figure S2.** Contribution of predators and prey to temporal changes in their interaction potentials, for all predator-prey pairs and in all local communities. The x-axis represents the temporal trend of predator relative abundances, and the y-axis represents the temporal trend of prey relative abundances. Pairs above the 1:1 line indicate increasing interaction potentials over time (i.e., positive temporal trends). Marginal density plots represent the distribution of temporal trends. Panel A (left): interactions recorded in the existing dataset; panel B (right): interactions predicted from the existing dataset. Only species pairs with statistically significant temporal trends of interaction potentials are included.

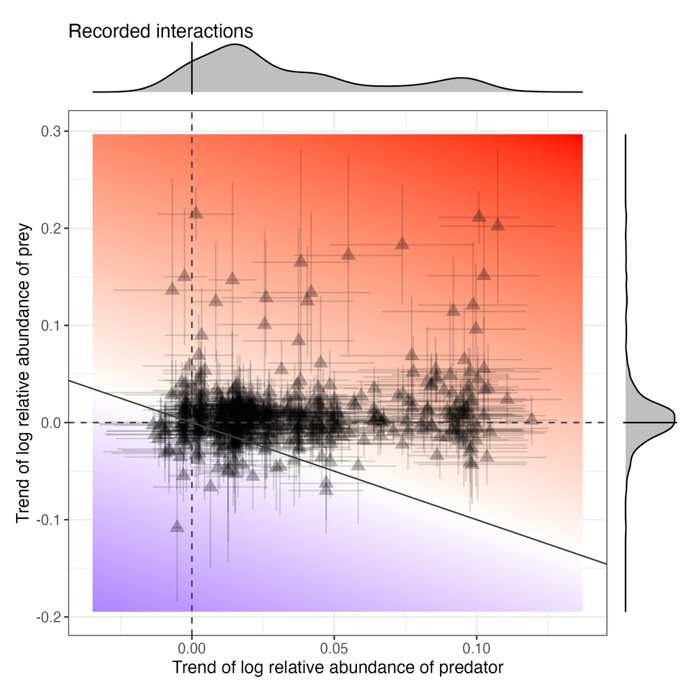

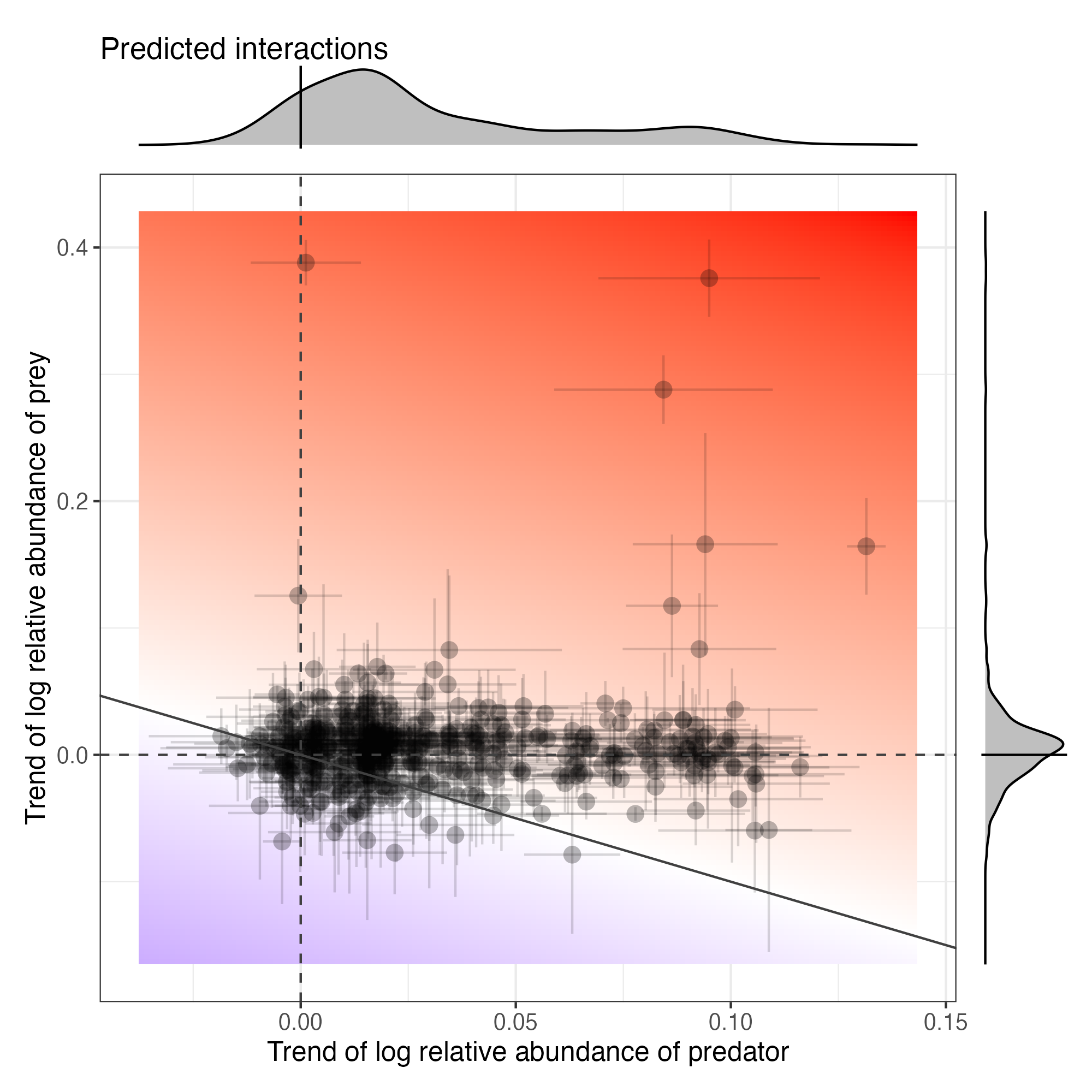

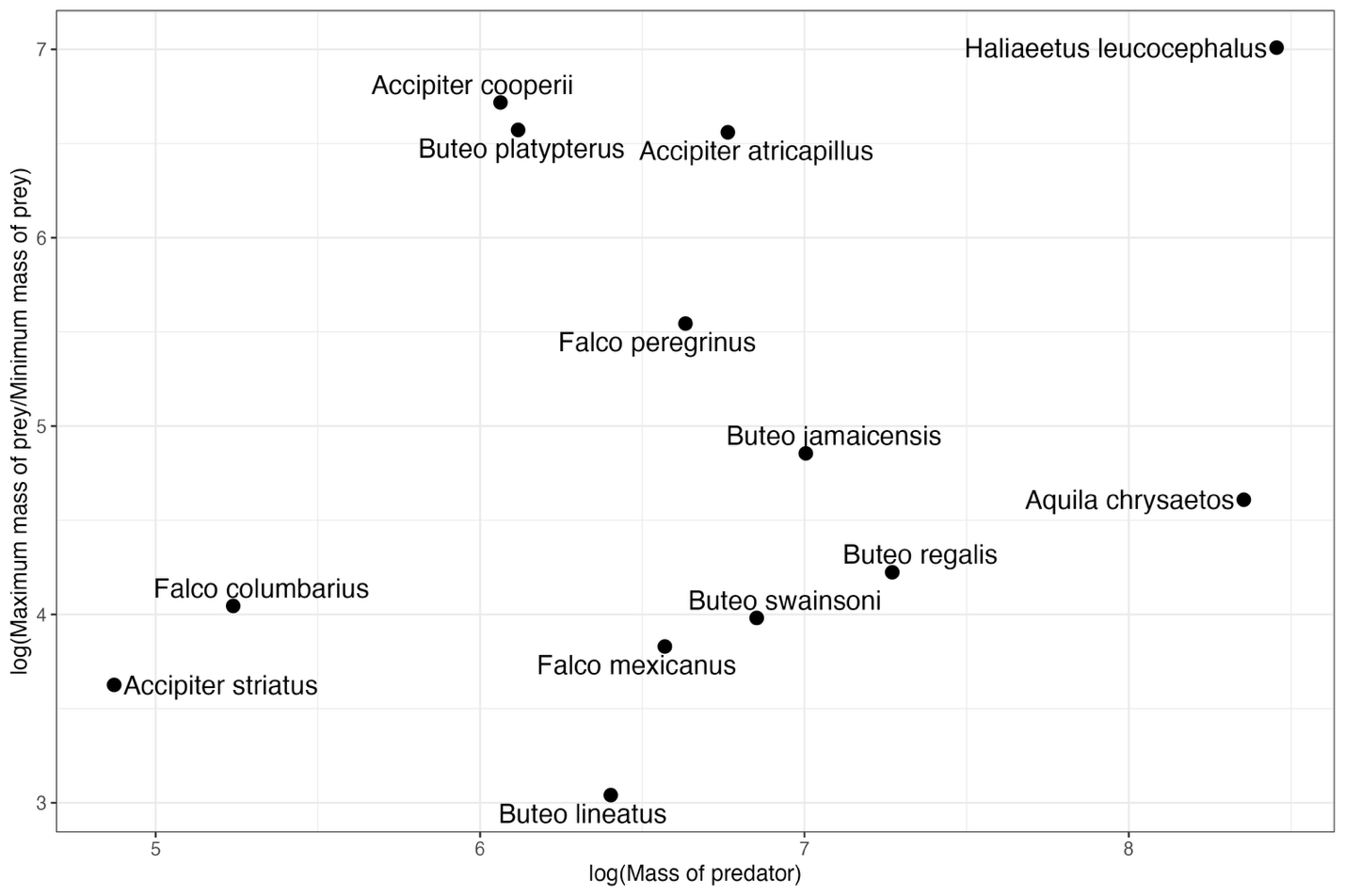
**Figure S3.** Ratio of maximum to minimum body mass, log-transformed, of recorded prey species for 14 predators included in the study. Larger ratios indicate that the predator species can prey on species with wider ranges of body mass.

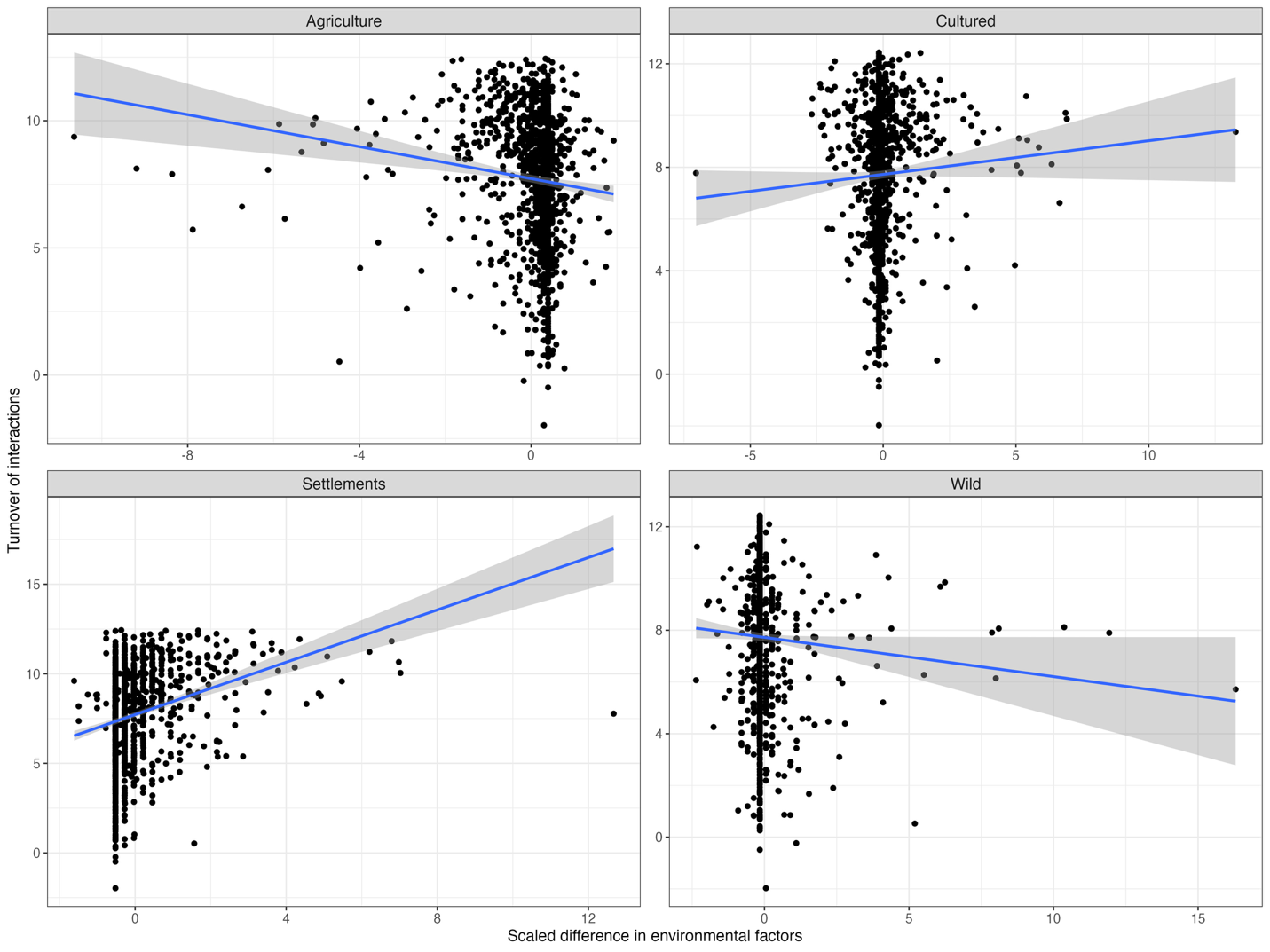
**Figure S4.** Associations between turnover of interaction networks in local communities (log-transformed; y-axis) and differences in four categories of human-induced land use changes (x-axis). The association with changes in wild land is the weakest.

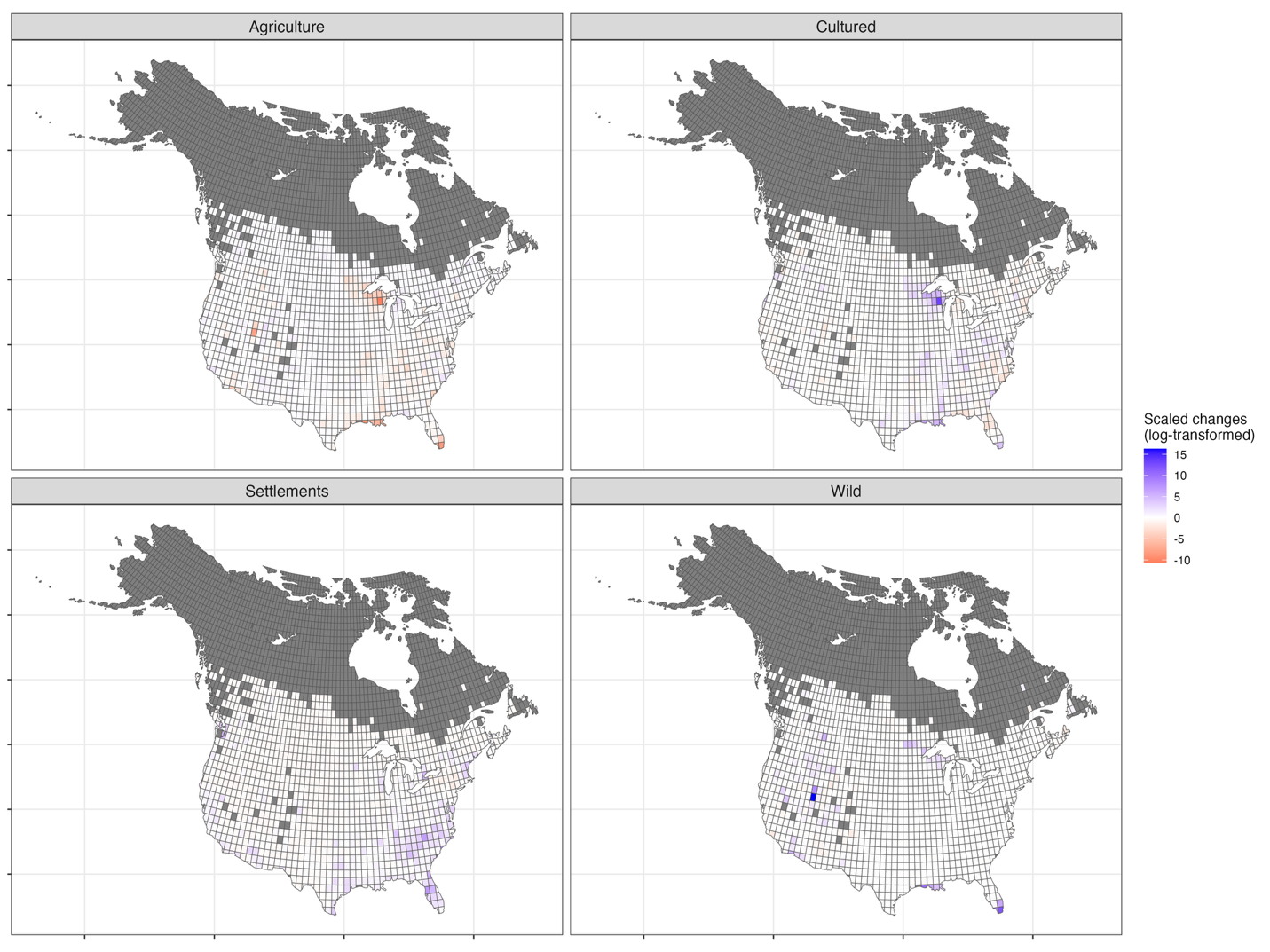

**Figure S5.** Maps of changes in four human-induced land use types between 1970 and 2021 in each one-degree latitude and longitude grid; see **Methods** for details of the calculations. These maps show the spatial variation in land use changes for different land types.

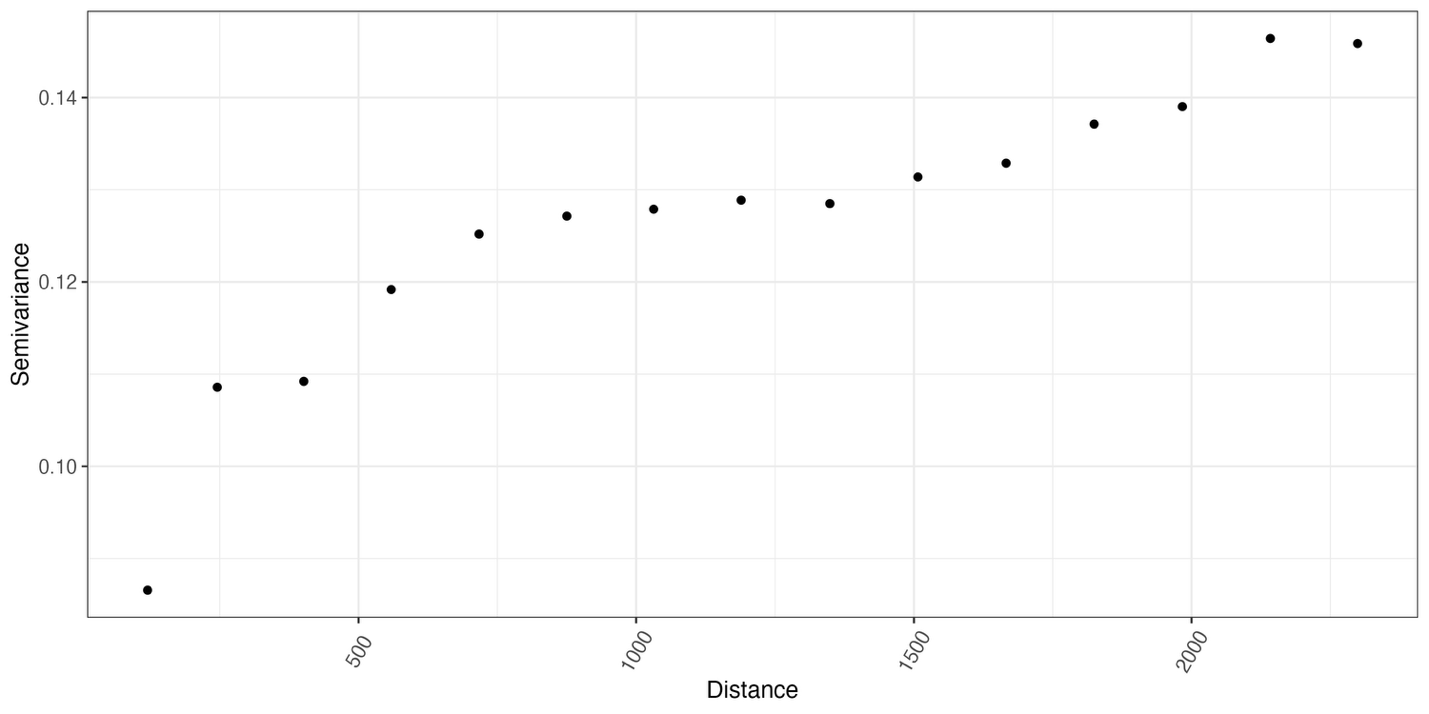
**Figure S6.** The variogram of the non-spatial linear model on turnover of interaction potentials between 1970 and 2021. See *Methods* for details of the model. The x-axis represents the distance, and the y-axis indicates the semivariance in the residuals of the model. The semivariance increases almost monotonically over distance, indicating a strong spatial signal in the residual of the non-spatial linear model that shows no sign of tapering off at a range of 2000km.

**Figure S7.** Most pairwise predator-prey interactions have increased their potential over time due to the increasing relative abundance of predators, not prey. These results are part of the sensitivity analysis, where we allowed for the threshold of functional compatibility to scale with the number of species in a community. Panel A (left): map showing the net count of increased interaction potentials, calculated by the number of pairs with increasing interaction potentials minus that of decreasing potentials in each one-degree latitude and longitude grid. Panel B (right): contribution of predators and prey to temporal changes in their interaction potentials, for all predator-prey pairs and in all local communities. For each pairwise interaction, the x-axis represents the mean temporal trend of predator relative abundances, and the y-axis represents the mean temporal trend of prey relative abundances, with associated standard deviations. Pairs above the 1:1 line indicate increasing interaction potentials over time (i.e., positive temporal trends). Marginal density plots represent the distribution of temporal trends. Both panels included interactions recorded in the existing dataset (triangles), and interactions predicted by the “functional compatibility” (dots). For both panels, one species pair will be counted multiple times if its interaction potential significantly increases/decreases in multiple local communities. Compare to Figure 1 where functional compatibility does not scale with the number of species in a community.

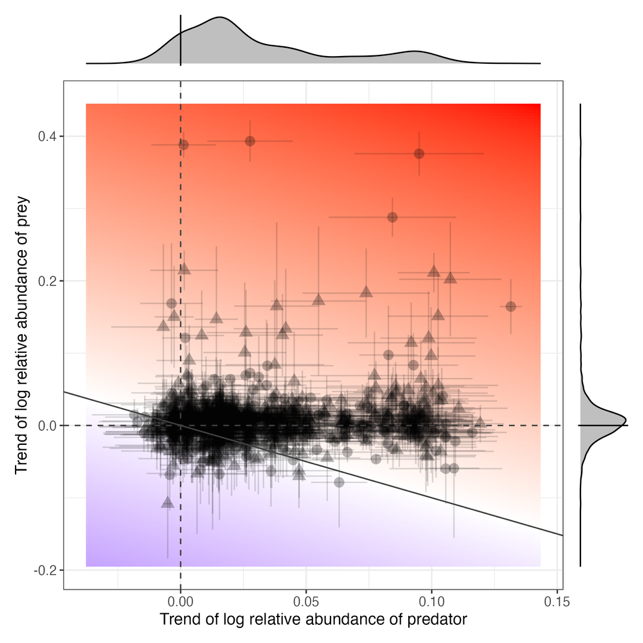

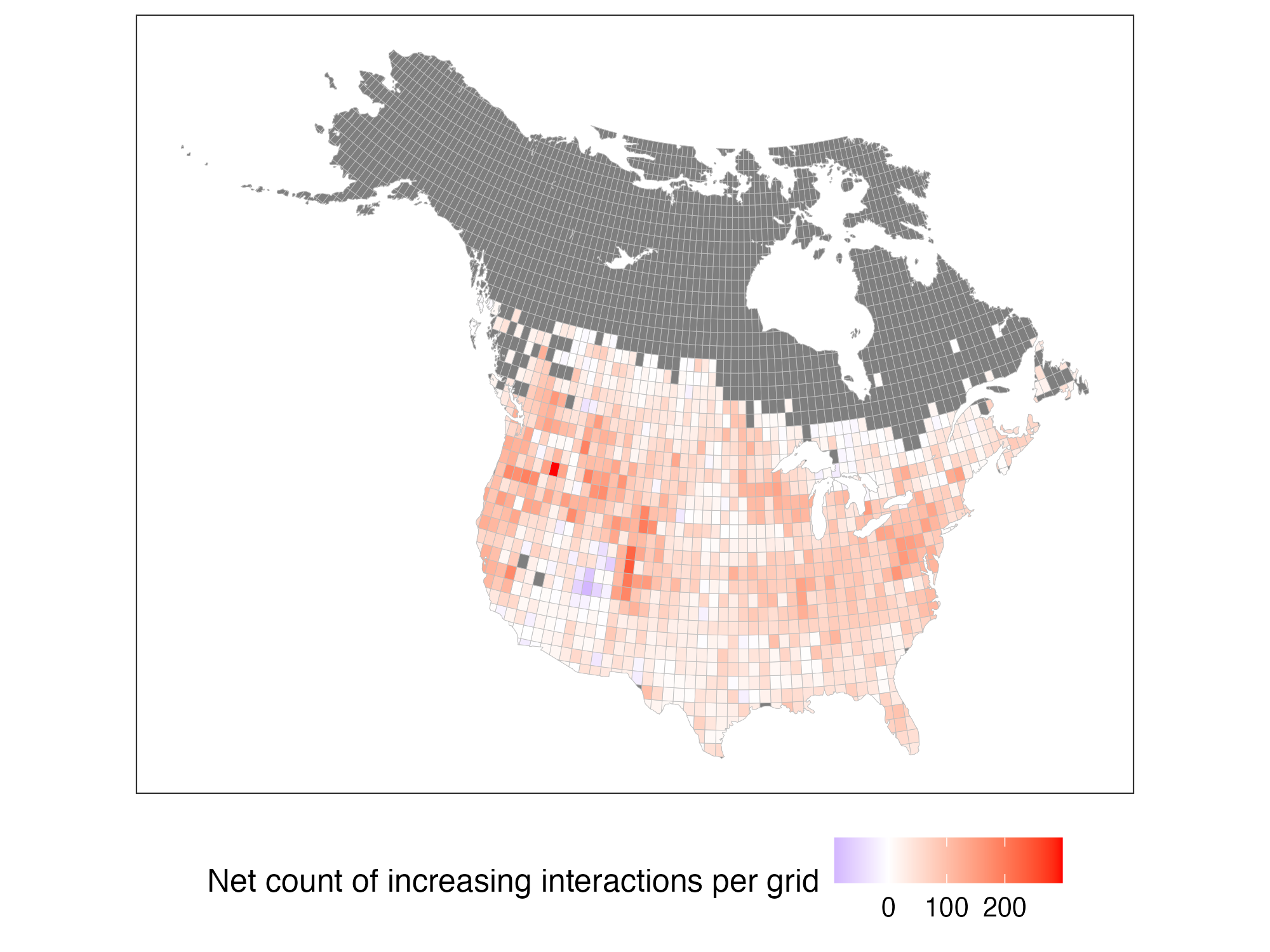

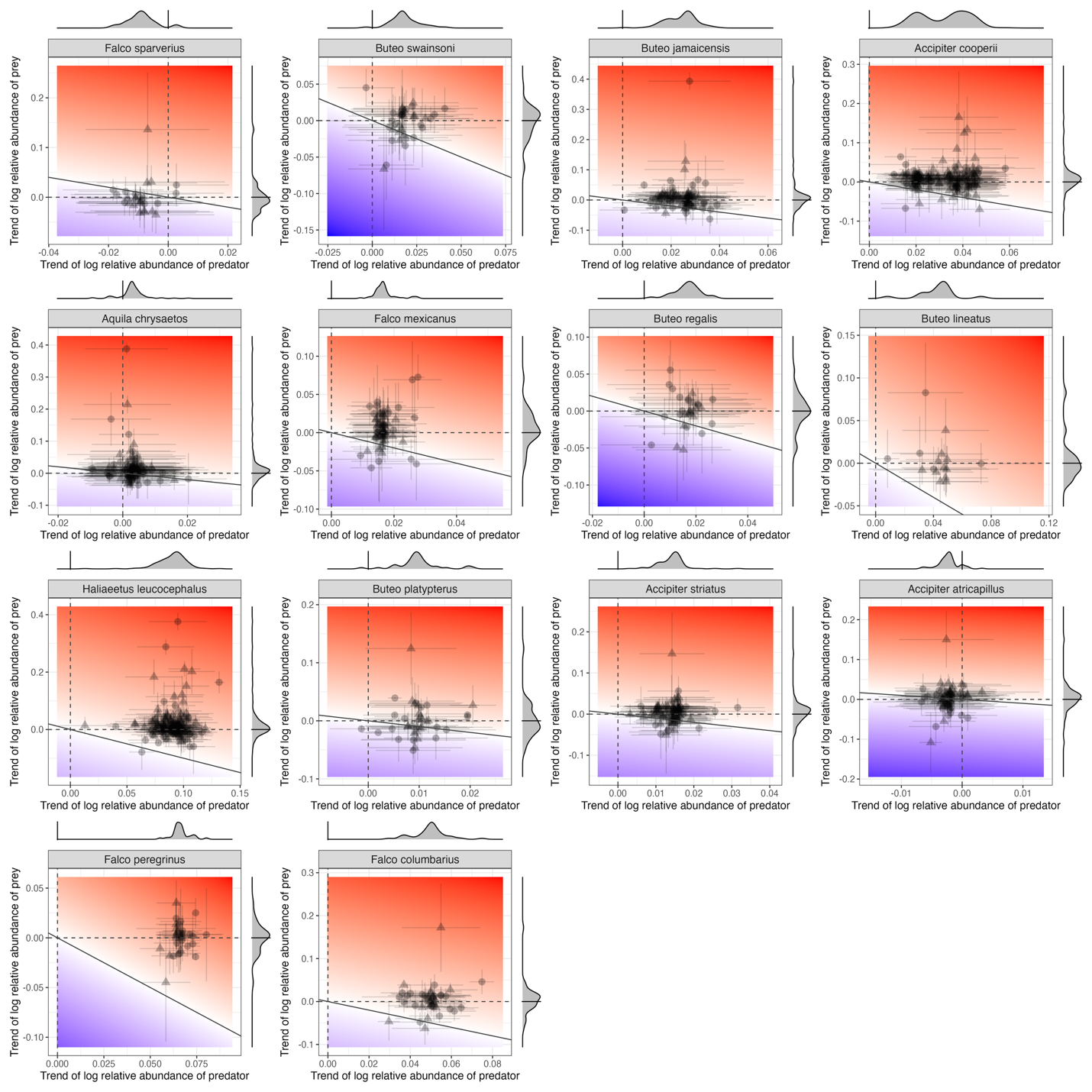

**Figure S8.** Contribution of predators and prey to temporal changes in their interaction potentials, for all pairs across all local communities by predator species. These results are part of the sensitivity analysis, where we allowed for the threshold of functional compatibility to scale with the number of species in a community. For each pairwise interaction, the x-axis represents the mean temporal trend of predator relative abundances, and the y-axis represents the mean temporal trend of prey relative abundances, with associated standard deviations. Pairs above the 1:1 line indicate increasing interaction potentials over time (i.e., positive temporal trends). Marginal density plots represent the distribution of temporal trends. All panels included interactions recorded in the existing dataset (triangles), and interactions predicted by the “functional compatibility” (dots). One species pair will be counted multiple times if its interaction potential significantly increases/decreases in multiple local communities. Compare to Figure 2 where functional compatibility does not scale with the number of species in a community.
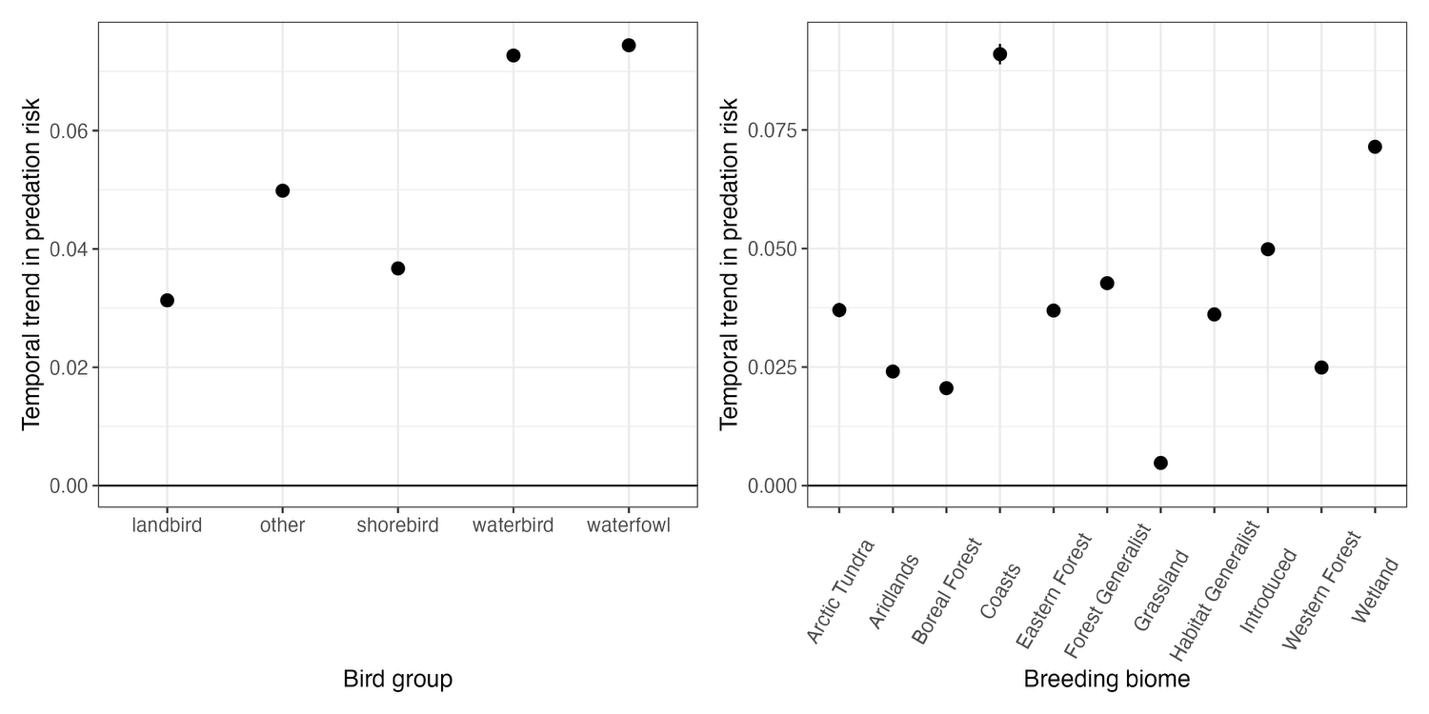

**Figure S9.** Temporal trends of overall predation risks by functional groups and breeding biomes of recorded and predicted prey species; points and lines indicate effect sizes and standard errors. These results are part of the sensitivity analysis, where we allowed for the threshold of functional compatibility to scale with the number of species in a community. Panel A (left): temporal trends by functional group; panel B (right): temporal trends by breeding biomes. Compare to Figure 3 where functional compatibility does not scale with the number of species in a community.

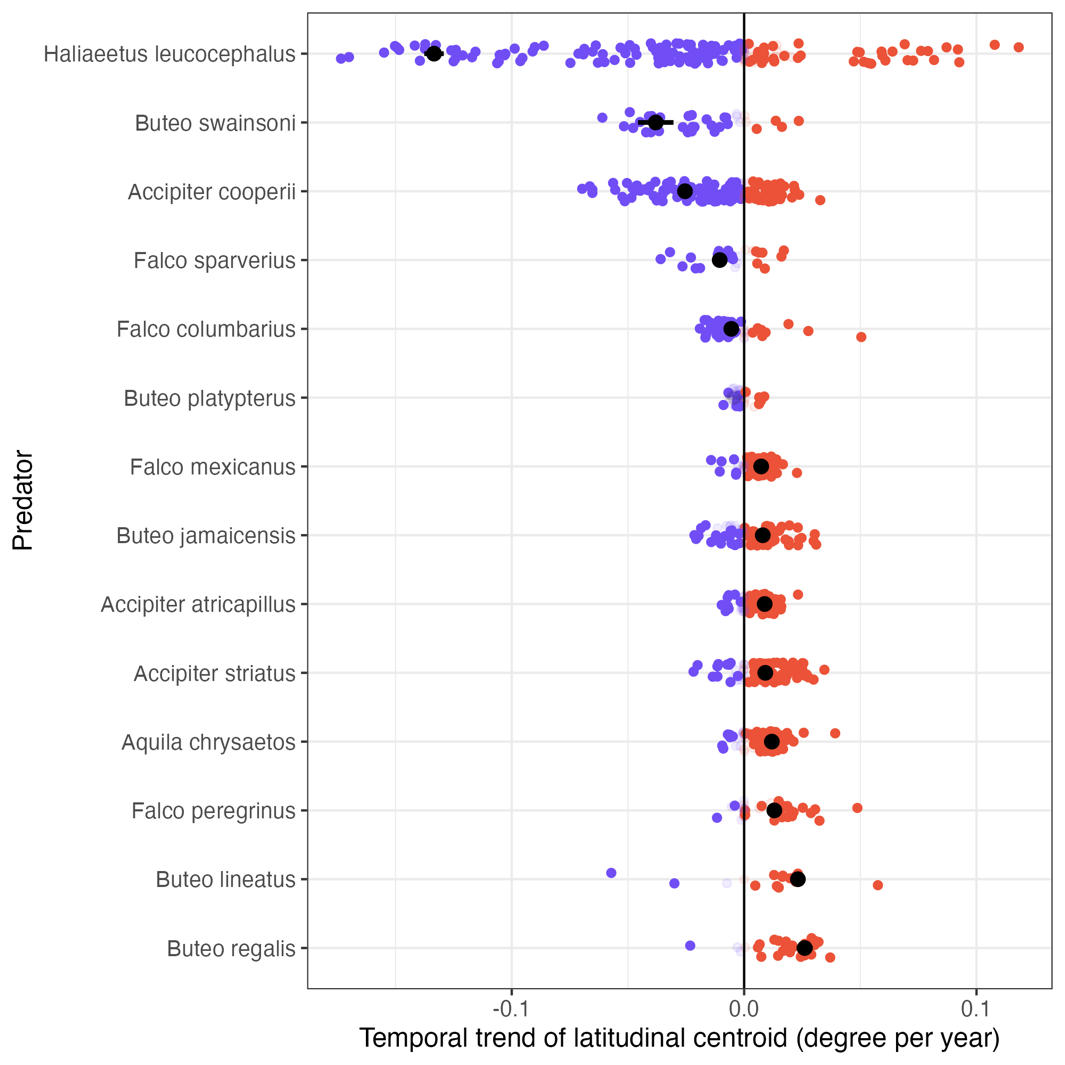

**Figure S10.** Latitudinal shifts over time of predator abundances in their entire ranges (black dots), and in their spatial overlap with their prey species (colored dots), including both recorded and predicted interactions. These results are part of the sensitivity analysis, where we allowed for the threshold of functional compatibility to scale with the number of species in a community. For a predator, because the spatial overlap with each prey species is different, each predator-prey pair will have an individual latitudinal shift. On the x-axis, positive numbers indicate northward shifts, negative numbers indicate southward shifts. Non-significant shifts are shown in lighter shades. Compare to Figure 4 where functional compatibility does not scale with the number of species in a community.

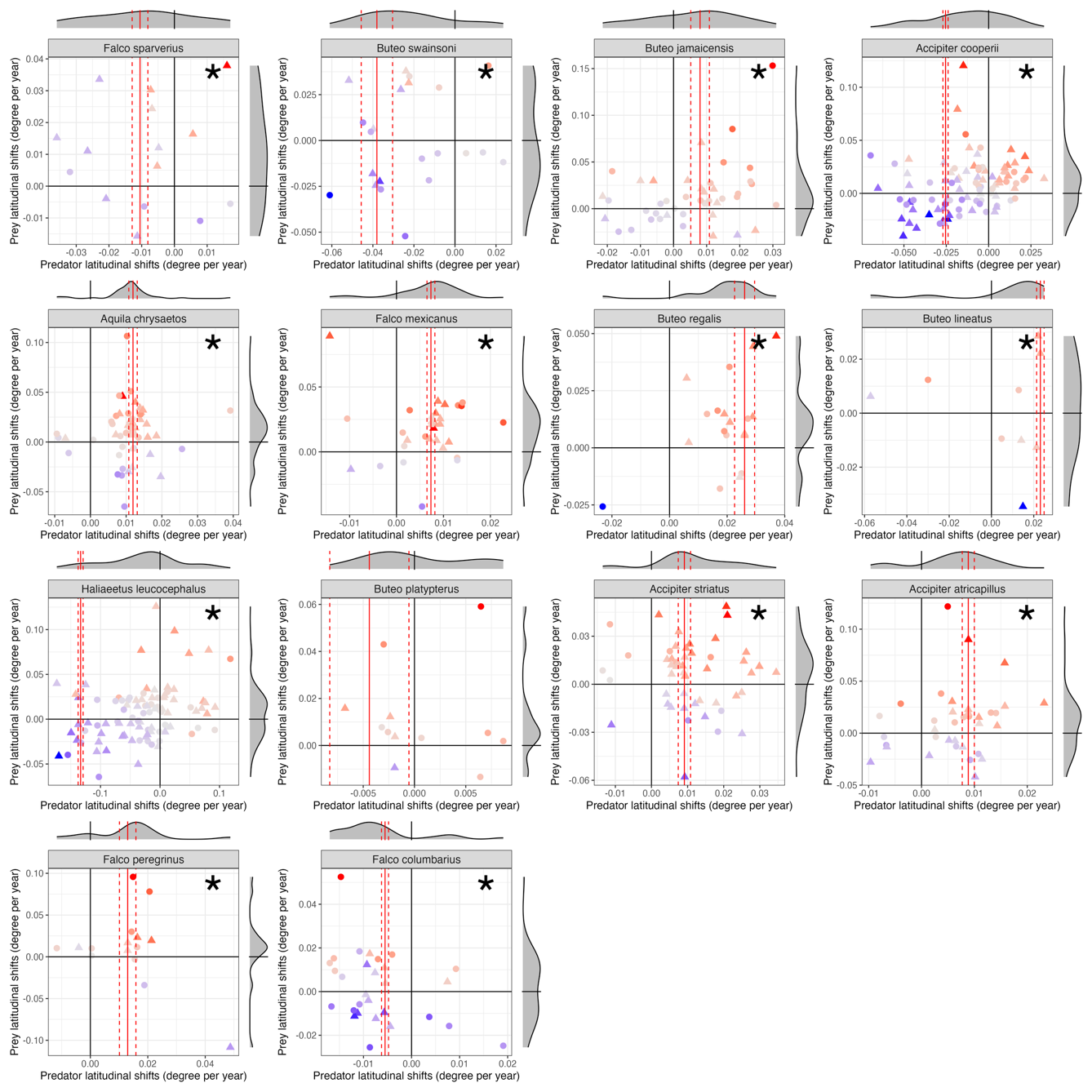

**Figure S11.** Latitudinal shifts of interaction potentials, decomposed by the latitudinal shifts of predators and prey abundances. Each panel represents both recorded and predicted interactions of one predator. These results are part of the sensitivity analysis, where we allowed for the threshold of functional compatibility to scale with the number of species in a community. The x- and y-axes indicate the latitudinal shifts of predators and prey, respectively, with positive values indicating northward shifts, and the marginal density plots show the distribution of latitudinal shifts, with black solid lines indicating 0 (no shift). Colors of each dot indicate the latitudinal shifts of each pairwise interaction potential (red: shifting significantly north; blue: shifting significantly south), dots indicate predicted interactions, and triangles indicate recorded interactions. The red vertical lines indicate the fitted slope and standard errors of the overall latitudinal shift of the predator, and the asterisks indicate that the means of the distributions of shifts in predator interaction potentials are significantly different from zero. Compare to Figure 5 where functional compatibility does not scale with the number of species in a community.

**Figure S12.** Most pairwise predator-prey interactions have increased their potential over time due to the increasing absolute abundance of predators, not prey. These results are part of the sensitivity analysis, where we calculated interaction potentials using absolute instead of relative abundances. Panel A (left): map showing the net count of increased interaction potentials, calculated by the number of pairs with increasing interaction potentials minus that of decreasing potentials in each one-degree latitude and longitude grid. Panel B (right): contribution of predators and prey to temporal changes in their interaction potentials, for all predator-prey pairs and in all local communities. For each pairwise interaction, the x-axis represents the mean temporal trend of predator relative abundances, and the y-axis represents the mean temporal trend of prey relative abundances, with associated standard deviations. Pairs above the 1:1 line indicate increasing interaction potentials over time (i.e., positive temporal trends). Marginal density plots represent the distribution of temporal trends. Both panels included interactions recorded in the existing dataset (triangles), and interactions predicted by the “functional compatibility” (dots). For both panels, one species pair will be counted multiple times if its interaction potential significantly increases/decreases in multiple local communities. Compare to Figure 1’s results with relative abundances.

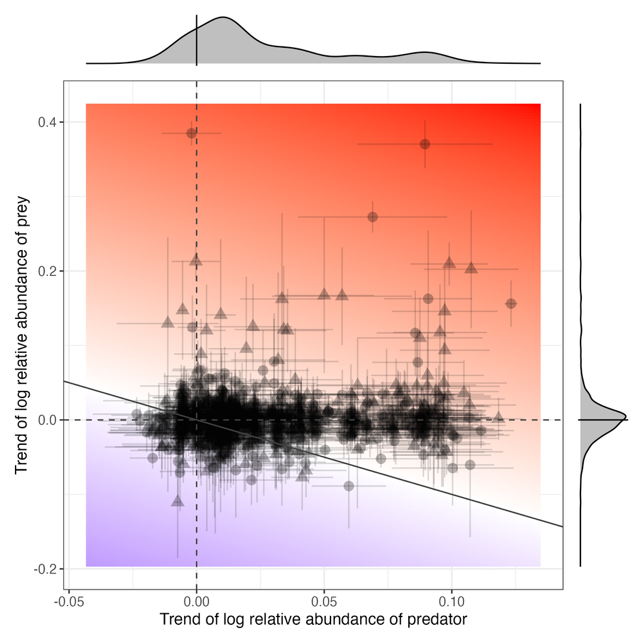

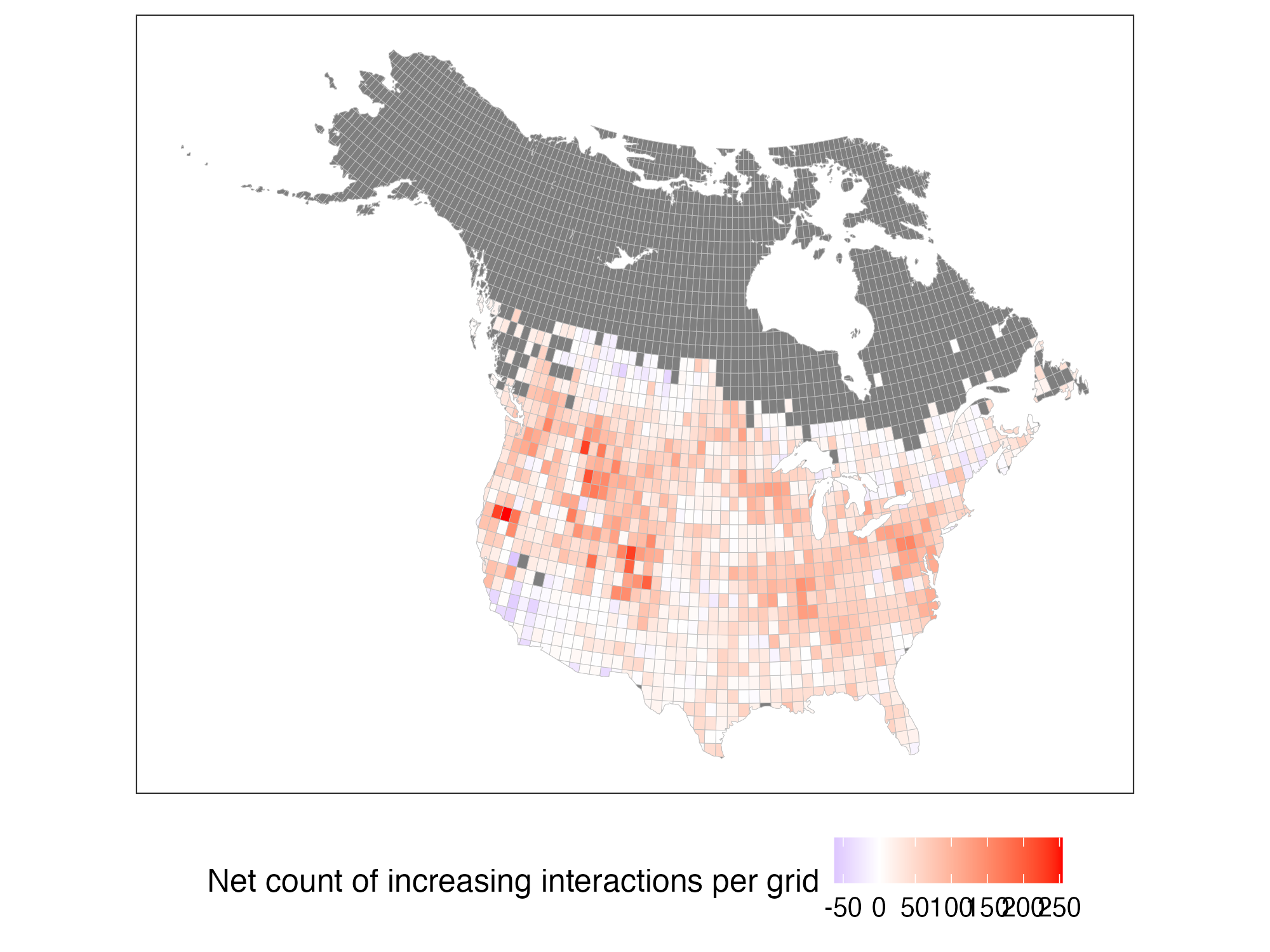

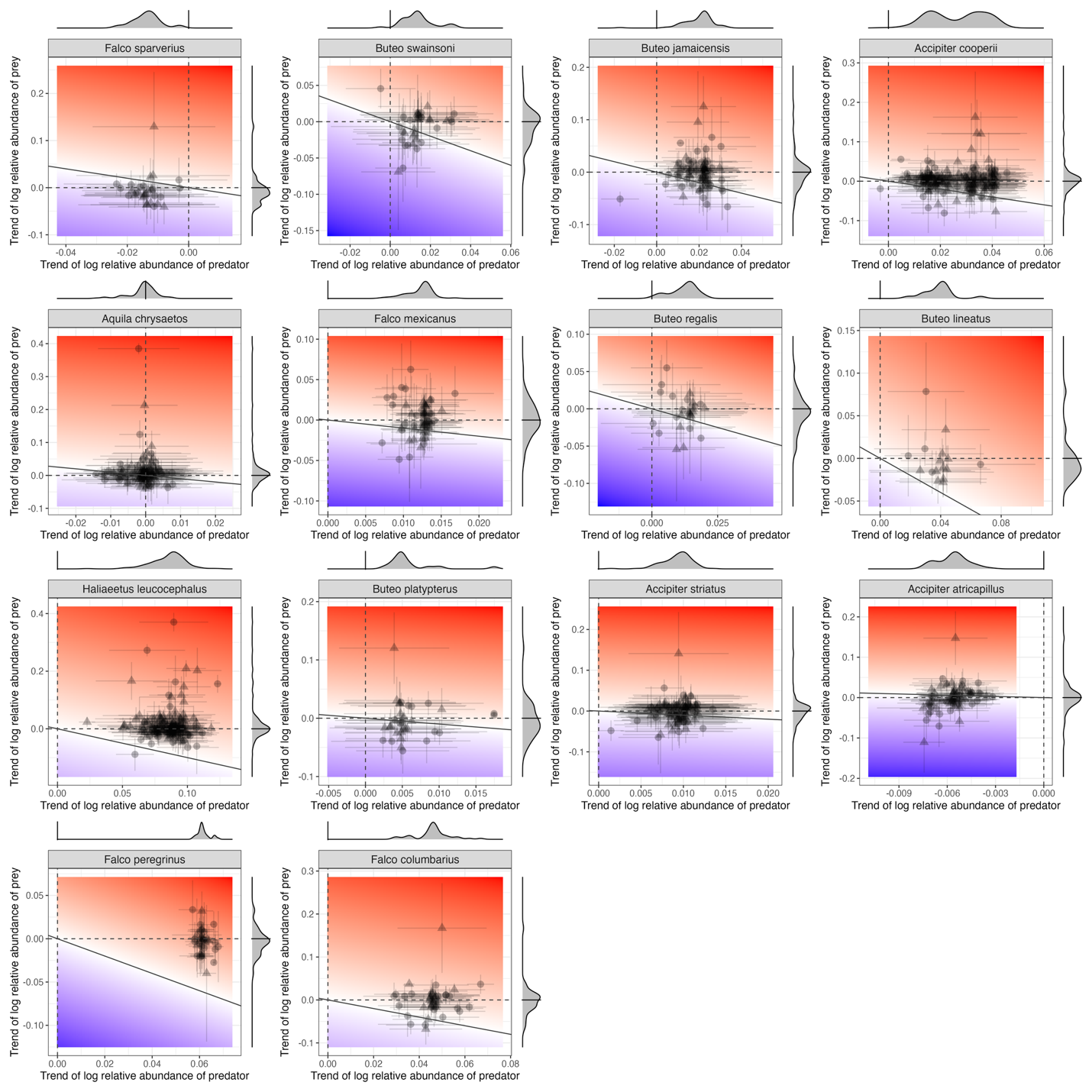

**Figure S13.** Contribution of predators and prey to temporal changes in their interaction potentials, for all pairs across all local communities by predator species. These results are part of the sensitivity analysis, where we calculated interaction potentials using absolute instead of relative abundances. For each pairwise interaction, the x-axis represents the mean temporal trend of predator relative abundances, and the y-axis represents the mean temporal trend of prey relative abundances, with associated standard deviations. Pairs above the 1:1 line indicate increasing interaction potentials over time (i.e., positive temporal trends). Marginal density plots represent the distribution of temporal trends. All panels included interactions recorded in the existing dataset (triangles), and interactions predicted by the “functional compatibility” (dots). One species pair will be counted multiple times if its interaction potential significantly increases/decreases in multiple local communities. Compare to Figure 2’s results with relative abundances

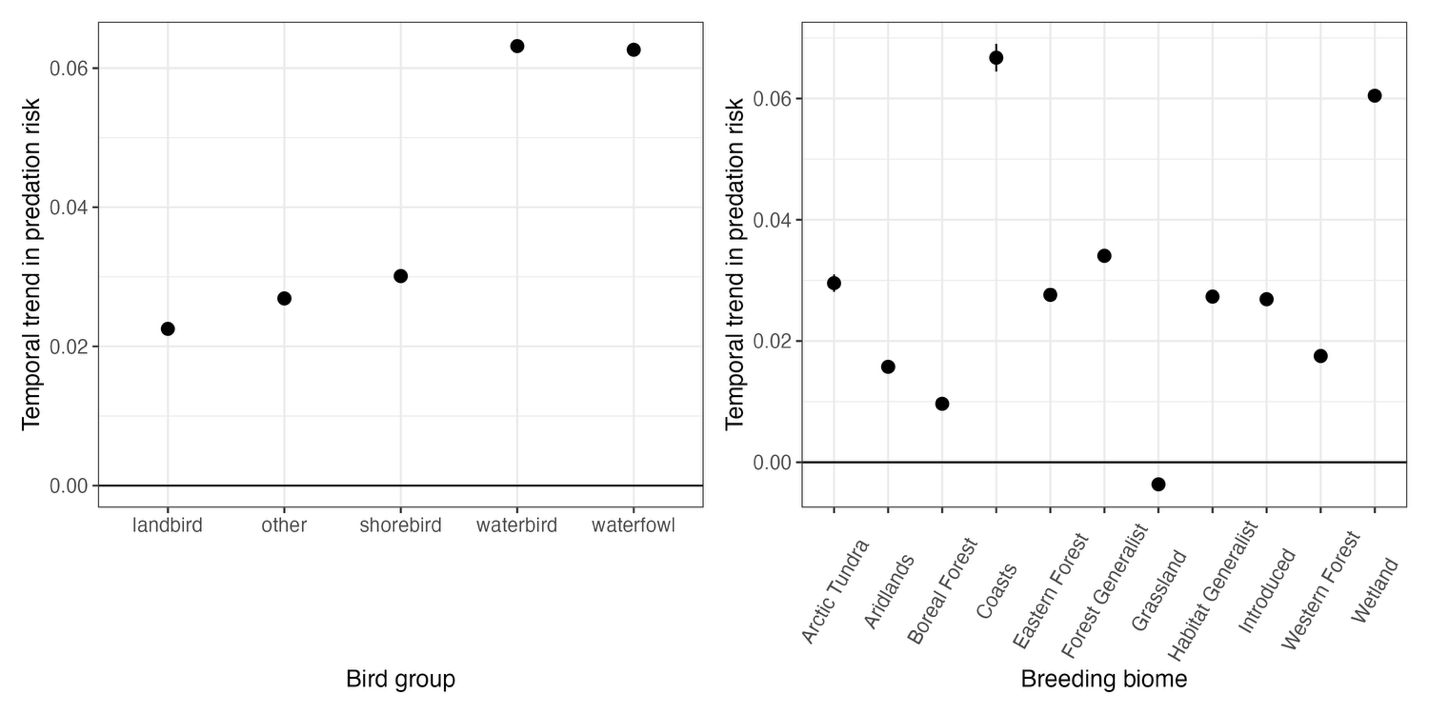
**Figure S14.** Temporal trends of overall predation risks by functional groups and breeding biomes of recorded and predicted prey species; points and lines indicate effect sizes and standard errors. These results are part of the sensitivity analysis, where we calculated interaction potentials using absolute instead of relative abundances. Panel A (left): temporal trends by functional group; panel B (right): temporal trends by breeding biomes. Note that grassland birds show decreasing predation risks over time when calculated using absolute abundances, possibly due to the strong decline in their populations over time. Compare to Figure 3’s results with relative abundances

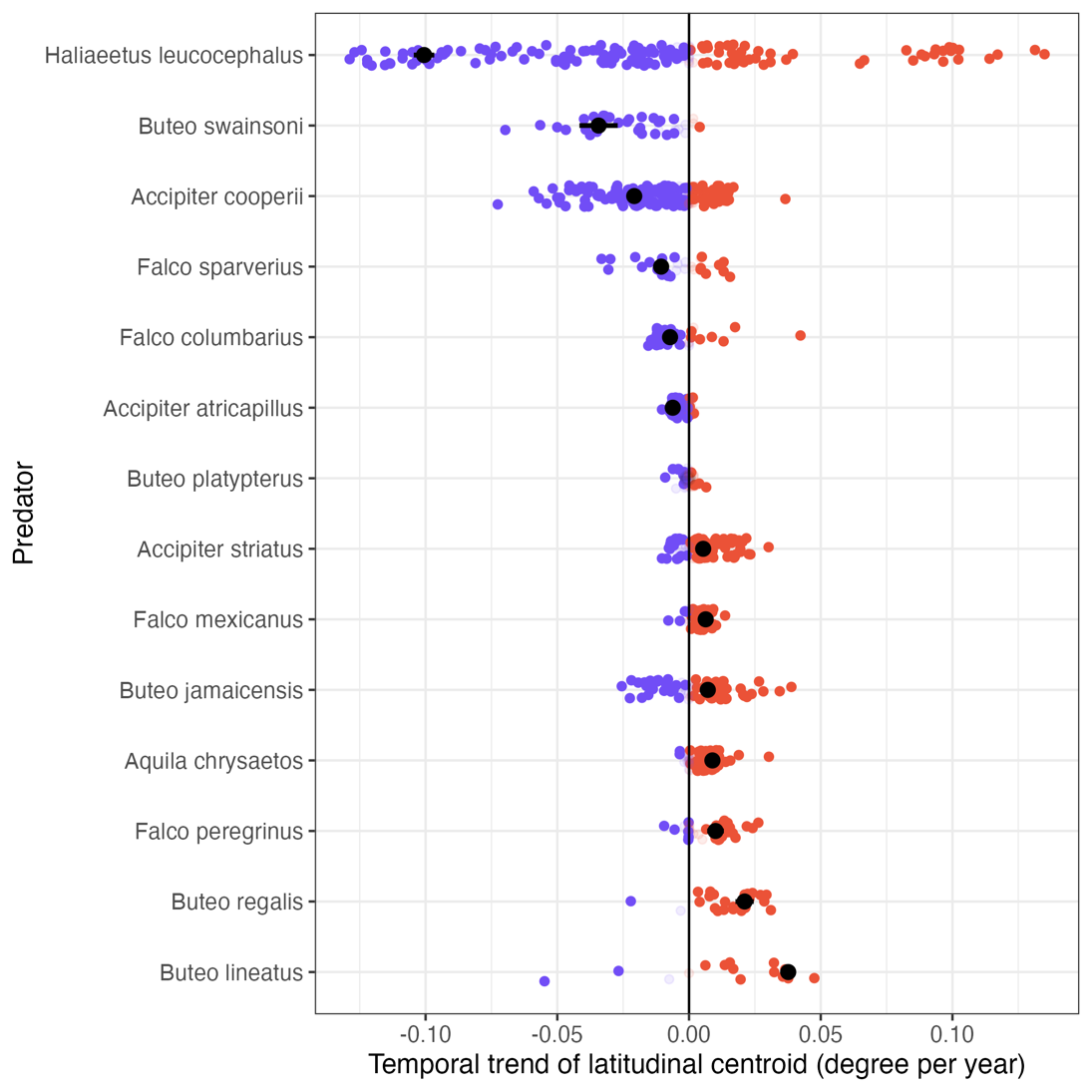

**Figure S15.** Latitudinal shifts over time of predator abundances in their entire ranges (black dots), and in their spatial overlap with their prey species (colored dots), including both recorded and predicted interactions. These results are part of the sensitivity analysis, where we calculated interaction potentials using absolute instead of relative abundances. For a predator, because the spatial overlap with each prey species is different, each predator-prey pair will have an individual latitudinal shift. On the x-axis, positive numbers indicate northward shifts, negative numbers indicate southward shifts. Non-significant shifts are shown in lighter shades. Compare to Figure 4’s results with relative abundances

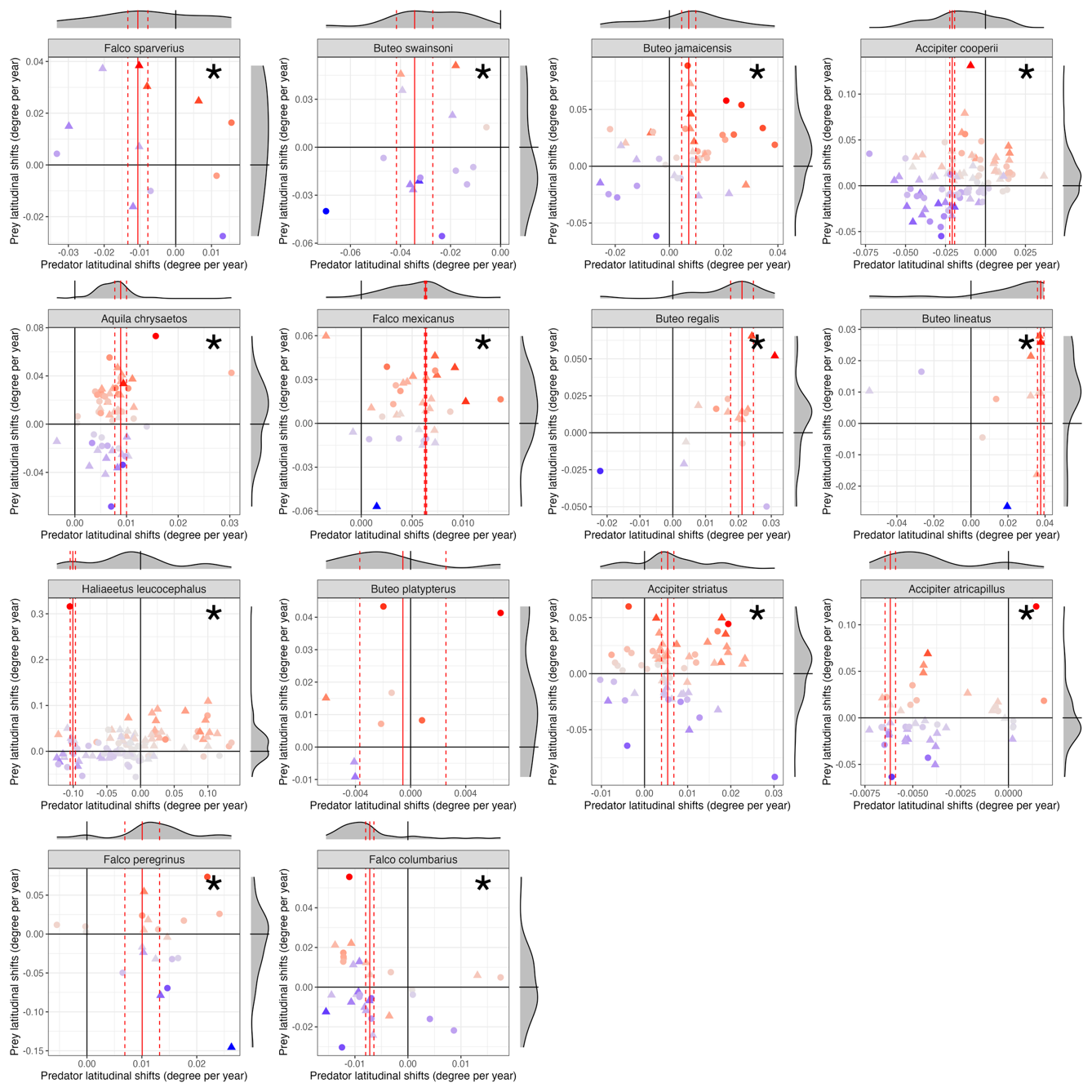
**Figure S16.** Latitudinal shifts of interaction potentials, decomposed by the latitudinal shifts of predators and prey abundances. Each panel represents both recorded and predicted interactions of one predator. These results are part of the sensitivity analysis, where we calculated interaction potentials using absolute instead of relative abundances. The x- and y-axes indicate the latitudinal shifts of predators and prey, respectively, with positive values indicating northward shifts, and the marginal density plots show the distribution of latitudinal shifts, with black solid lines indicating 0 (no shift). Colors of each dot indicate the latitudinal shifts of each pairwise interaction potential (red: shifting significantly north; blue: shifting significantly south), dots indicate predicted interactions, and triangles indicate recorded interactions. The red vertical lines indicate the fitted slope and standard errors of the overall latitudinal shift of the predator, and the asterisks indicate that the means of the distributions of shifts in predator interaction potentials are significantly different from zero. Compare to Figure 5’s results with relative abundances.

**Figure S17.** Most pairwise predator-prey interactions have increased their potential over time due to the increasing abundance of predators, not prey. Panel A (left): map showing the net count of increased interaction potentials, calculated by the number of pairs with increasing interaction potentials minus that of decreasing potentials in each one-degree latitude and longitude grid, including pairs that show both significant and non-significant temporal trends. Panel B (right): contribution of predators and prey to temporal changes in their interaction potentials, for predator-prey pairs showing non-significant temporal trends only. For each pairwise interaction, the x-axis represents the mean temporal trend of predator relative abundances, and the y-axis represents the mean temporal trend of prey relative abundances, with associated standard deviations. Pairs above the 1:1 line indicate increasing interaction potentials over time (i.e., positive temporal trends). Marginal density plots represent the distribution of temporal trends. Both panels included interactions recorded in the existing dataset (triangles) and interactions predicted by the “functional compatibility” (dots). For both panels, one species pair will be counted multiple times if its interaction potential significantly increases/decreases in multiple local communities. Compare to Figure 1’s results with interaction pairs with significant temporal trends only.

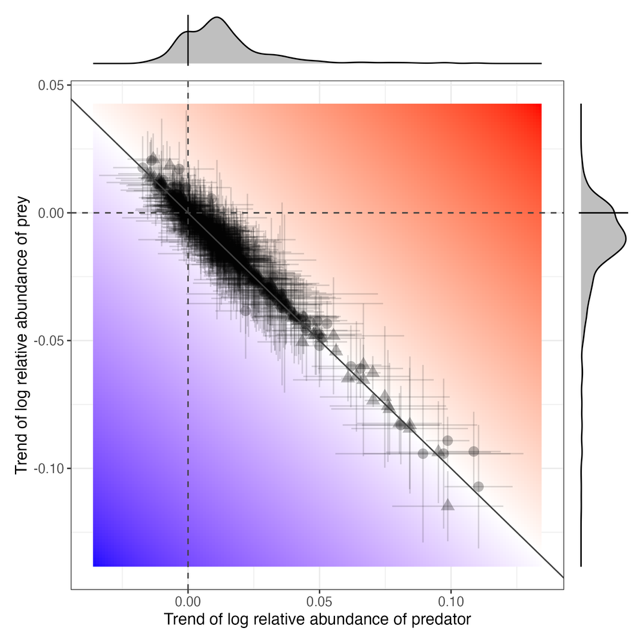

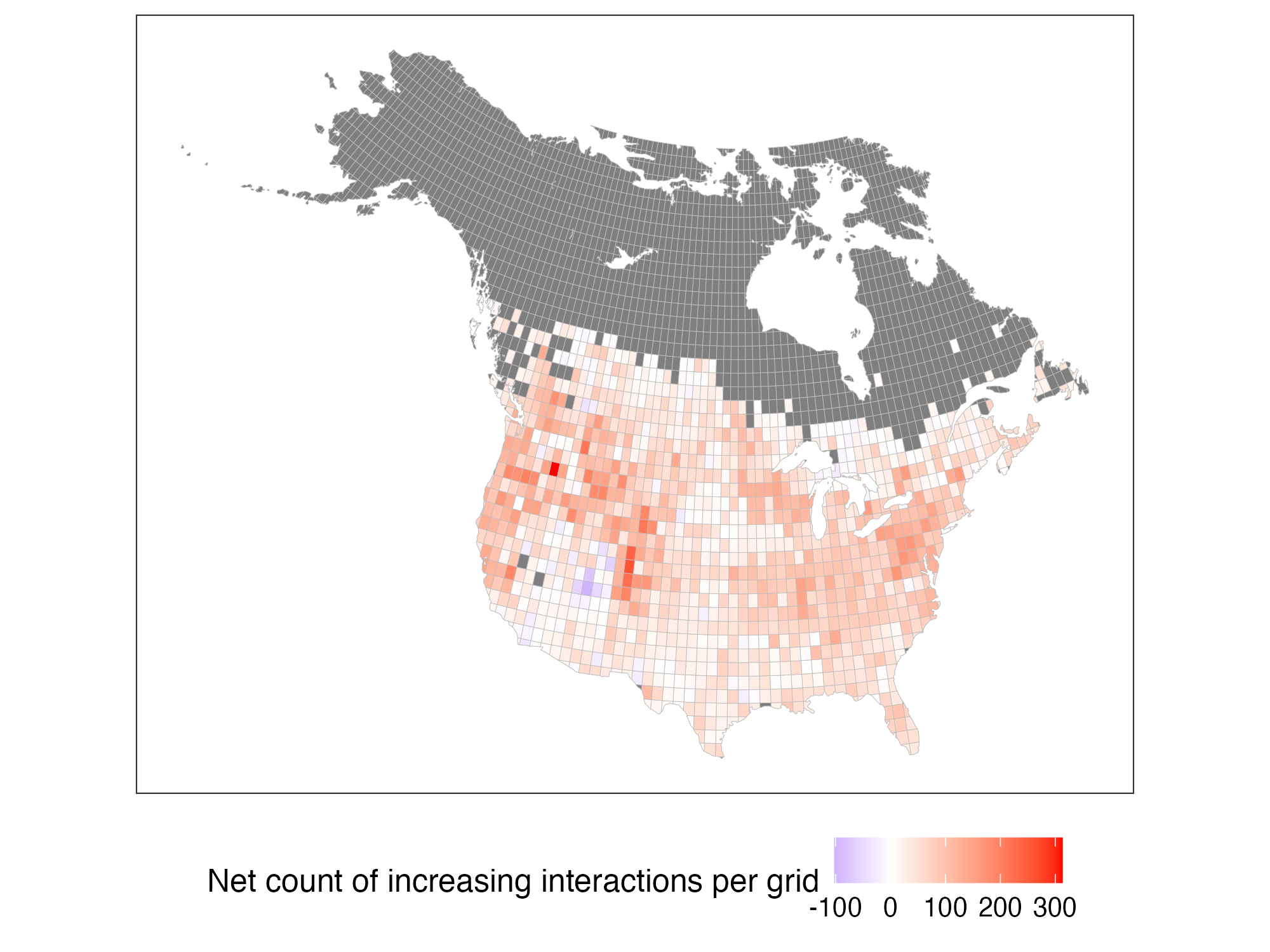

**Table S1.** Statistics of linear mixed effects models for the temporal trends of predation risks for prey species by groups. The upper table shows the results by functional groups, lower table shows the results by breeding biomes.

| **Parameters** | **Estimate** | **Std. Error** | **df** | **t value** | **Adjusted *p* value** | **Group** |
| --- | --- | --- | --- | --- | --- | --- |
| (Intercept) | -15.811829 | 0.1319793 | 271.197206 | -119.80537 | **<0.00001** | landbird |
| year_since_1969 | 0.03147079 | 8.24E-05 | 2773348.23 | 382.121203 | **<0.00001** | landbird |
| (Intercept) | -19.49082 | 0.37330267 | 12.679676 | -52.211841 | **<0.00001** | shorebird |
| year_since_1969 | 0.03815552 | 3.35E-04 | 92543.4978 | 113.990421 | **<0.00001** | shorebird |
| (Intercept) | -21.475931 | 0.2459613 | 44.991039 | -87.31427 | **<0.00001** | waterbird |
| year_since_1969 | 0.07240634 | 3.28E-04 | 158603.847 | 220.537286 | **<0.00001** | waterbird |
| (Intercept) | -19.99579 | 0.36763656 | 28.6950331 | -54.390103 | **<0.00001** | waterfowl |
| year_since_1969 | 0.07022692 | 2.44E-04 | 222496.518 | 287.645469 | **<0.00001** | waterfowl |
| (Intercept) | -17.230321 | 1.64156718 | 6.98113706 | -10.496263 | **<0.00001** | other |
| year_since_1969 | 0.03624844 | 3.18E-04 | 212109.507 | 113.946304 | **<0.00001** | other |

| **parameters** | **Estimate** | **Std. Error** | **df** | **t value** | **Adjusted *p* value** | **Breeding biome** |
| --- | --- | --- | --- | --- | --- | --- |
| (Intercept) | -16.414673 | 0.26519014 | 44.0135452 | -61.897751 | **<0.00001** | Eastern Forest |
| year_since_1969 | 0.03742953 | 1.55E-04 | 549790.499 | 242.096253 | **<0.00001** | Eastern Forest |
| (Intercept) | -16.599323 | 0.37867722 | 42.4804646 | -43.835019 | **<0.00001** | Aridlands |
| year_since_1969 | 0.02390003 | 3.03E-04 | 141321.107 | 78.9966579 | **<0.00001** | Aridlands |
| (Intercept) | -16.132047 | 0.22620762 | 62.0706469 | -71.315223 | **<0.00001** | Western Forest |
| year_since_1969 | 0.02506117 | 2.26E-04 | 220347.243 | 111.090013 | **<0.00001** | Western Forest |
| (Intercept) | -16.388839 | 0.43012669 | 29.3603395 | -38.102354 | **<0.00001** | Boreal Forest |
| year_since_1969 | 0.01920441 | 2.46E-04 | 160769.955 | 78.2086415 | **<0.00001** | Boreal Forest |
| (Intercept) | -20.96377 | 0.20990683 | 84.5442781 | -99.871784 | **<0.00001** | Wetland |
| year_since_1969 | 0.06884089 | 1.86E-04 | 459346.81 | 369.23315 | **<0.00001** | Wetland |
| (Intercept) | -16.229741 | 0.61039783 | 22.2424569 | -26.588791 | **<0.00001** | Habitat Generalist |
| year_since_1969 | 0.03619846 | 1.76E-04 | 601151.166 | 205.384232 | **<0.00001** | Habitat Generalist |
| (Intercept) | -17.702427 | 0.5741062 | 7.8736301 | -30.83476 | **<0.00001** | Arctic Tundra |
| year_since_1969 | 0.03702951 | 0.00152967 | 1695.1187 | 24.2075876 | **<0.00001** | Arctic Tundra |
| (Intercept) | -16.674112 | 0.36419652 | 35.687542 | -45.783282 | **<0.00001** | Forest Generalist |
| year_since_1969 | 0.04325518 | 1.36E-04 | 691288.292 | 318.294069 | **<0.00001** | Forest Generalist |
| (Intercept) | -21.106998 | 1.04333342 | 3.77053432 | -20.230347 | **<0.00001** | Coasts |
| year_since_1969 | 0.09100524 | 0.00216601 | 1534.90013 | 42.0151681 | **<0.00001** | Coasts |
| (Intercept) | -15.325279 | 0.49943613 | 24.8763874 | -30.685164 | **<0.00001** | Grassland |
| year_since_1969 | 0.00435323 | 2.07E-04 | 416282.393 | 21.0030585 | 0.16057 | Grassland |
| (Intercept) | -17.230321 | 1.64156718 | 6.98113706 | -10.496263 | **<0.00001** | Introduced |
| year_since_1969 | 0.03624844 | 3.18E-04 | 212109.507 | 113.946304 | **<0.00001** | Introduced |

**Table S2.** Statistics for the latitudinal shifts of each predator, estimated from linear models between the latitudinal centroid and time (see *Methods*).

| **Estimate** | **SE** | **statistic** | **Predator** | **Adjusted *p* value** |
| --- | --- | --- | --- | --- |
| -0.0105081 | 0.00239671 | -4.3843762 | *Falco sparverius* | **0.000102** |
| -0.0380354 | 0.00763398 | -4.9823748 | *Buteo swainsoni* | **0.0000145** |
| 0.00801418 | 0.0028087 | 2.85333823 | *Buteo jamaicensis* | **0.00848** |
| -0.0254503 | 0.00153488 | -16.581306 | *Accipiter cooperii* | **<0.00001** |
| 0.01192226 | 0.0011822 | 10.0848379 | *Aquila chrysaetos* | **<0.00001** |
| 0.0073343 | 8.38E-04 | 8.74936822 | *Falco mexicanus* | **<0.00001** |
| 0.02605704 | 0.00347729 | 7.49349361 | *Buteo regalis* | **<0.00001** |
| 0.02313013 | 0.00176773 | 13.0846676 | *Buteo lineatus* | **<0.00001** |
| -0.1334401 | 0.00419295 | -31.824898 | *Haliaeetus leucocephalus* | **<0.00001** |
| -0.0044008 | 0.00386101 | -1.1398061 | *Buteo platypterus* | 0.291 |
| 0.00911015 | 0.00170047 | 5.35742392 | *Accipiter striatus* | **<0.00001** |
| 0.00884313 | 0.00114396 | 7.7302572 | *Accipiter atricapillus* | **<0.00001** |
| 0.01299551 | 0.00287001 | 4.52803031 | *Falco peregrinus* | **0.0000656** |
| -0.0055129 | 7.33E-04 | -7.5221904 | *Falco columbarius* | **<0.00001** |

**Table S3.** Statistics of the linear model with spatial lag between turnover of predator-prey interactions and environmental changes between the years 1970 and 2021. Turnover was calculated as the difference between matrices of interaction potentials, including recorded predator-prey interactions only. Environmental change includes changes in land use, mean annual temperature, and mean annual precipitation, and the interaction between changes in temperature and precipitation within each local community. See *Methods* for details on model fitting. Compare to Table 1’s results with both recorded and predicted interactions.

| **Environmental change** | **Estimate** | **SE** | ***z* value** | ***p* value** |
| --- | --- | --- | --- | --- |
| (Intercept) | 0.71943 | 0.16849 | 4.26978 | **0.00002** |
| Temperature | 0.03778 | 0.05962 | 0.63377 | 0.52623 |
| Precipitation | 0.00287 | 0.07015 | 0.04088 | 0.96739 |
| Settlements | 0.35137 | 0.07956 | 4.41617 | **0.00001** |
| Cultured | 0.08890 | 0.12783 | 0.69540 | 0.48680 |
| Agriculture | 0.09862 | 0.12888 | 0.76523 | 0.44414 |
| Temperature: precipitation | 0.05110 | 0.06220 | 0.82164 | 0.41128 |
| N = 1179, df = 1170; $\rho$ = 0.899, *p* value for $\rho$ < 0.0001; pseudo R^2^ = 0.502 | | | | |

**Table S4.** Statistics of the linear model with spatial lag between turnover of predator-prey interactions and environmental changes between the years 1970 and 2021. These results are part of the sensitivity analysis, where we allowed for the threshold of functional compatibility to scale with the number of species in a community. Turnover was calculated as the difference between matrices of interaction potentials, including both recorded and predicted predator-prey interactions. Environmental change includes changes in land use, mean annual temperature, and mean annual precipitation, and the interaction between changes in temperature and precipitation within each local community. See *Methods* for details on model fitting. Compare to Table 1, where functional compatibility does not scale with the number of species in a community.

| **Environmental change** | **Estimate** | **SE** | ***z* value** | ***p* value** |
| --- | --- | --- | --- | --- |
| (Intercept) | 0.98611 | 0.21760 | 4.53184 | **0.00001** |
| Temperature | -0.00042 | 0.06300 | -0.00665 | 0.99470 |
| Precipitation | 0.03339 | 0.07397 | 0.45133 | 0.65175 |
| Settlements | 0.22719 | 0.08352 | 2.72008 | **0.00653** |
| Cultured | -0.09043 | 0.13464 | -0.67164 | 0.50181 |
| Agriculture | -0.09433 | 0.13578 | -0.69471 | 0.48724 |
| Temperature: precipitation | 0.03651 | 0.06546 | 0.55784 | 0.57695 |
| N = 1178, df = 1169; $\rho$ = 0.880, *p* value for $\rho$ < 0.0001; pseudo R^2^ = 0.428 | | | | |

**Table S5.** Statistics of linear mixed effects models for the temporal trends of predation risks for prey species by groups. These results are part of the sensitivity analysis, where we allowed for the threshold of functional compatibility to scale with the number of species in a community. The upper table shows the results by functional groups, lower table shows the results by breeding biomes. Compare to Table S1, where functional compatibility does not scale with the number of species in a community.

| **Parameters** | **Estimate** | **Std. Error** | **df** | **t value** | **Adjusted *p* value** | **Group** |
| --- | --- | --- | --- | --- | --- | --- |
| (Intercept) | -15.830915 | 0.13363817 | 266.328219 | -118.46103 | **<0.00001** | landbird |
| year_since_1969 | 0.03131754 | 8.46E-05 | 2669727.97 | 370.052871 | **<0.00001** | landbird |
| (Intercept) | -19.424799 | 0.37105805 | 12.6800935 | -52.349759 | **<0.00001** | shorebird |
| year_since_1969 | 0.03671585 | 3.39E-04 | 88906.4721 | 108.300823 | **<0.00001** | shorebird |
| (Intercept) | -21.431799 | 0.24456654 | 43.9995011 | -87.63177 | **<0.00001** | waterbird |
| year_since_1969 | 0.07271737 | 3.30E-04 | 154965.98 | 220.224228 | **<0.00001** | waterbird |
| (Intercept) | -20.276204 | 0.3749187 | 28.6491383 | -54.081604 | **<0.00001** | waterfowl |
| year_since_1969 | 0.07443062 | 2.51E-04 | 213656.781 | 297.059297 | **<0.00001** | waterfowl |
| (Intercept) | -17.350215 | 1.50982733 | 7.08618937 | -11.491523 | **<0.00001** | other |
| year_since_1969 | 0.04985819 | 3.54E-04 | 219802.117 | 140.722753 | **<0.00001** | other |

| **parameters** | **Estimate** | **Std. Error** | **df** | **t value** | **Adjusted *p* value** | **Breeding biome** |
| --- | --- | --- | --- | --- | --- | --- |
| (Intercept) | -16.510596 | 0.28612796 | 40.6077491 | -57.703537 | **<0.00001** | Eastern Forest |
| year_since_1969 | 0.03692613 | 1.63E-04 | 482979.327 | 226.277083 | **<0.00001** | Eastern Forest |
| (Intercept) | -16.573448 | 0.35487305 | 43.4531636 | -46.702469 | **<0.00001** | Aridlands |
| year_since_1969 | 0.02406847 | 3.10E-04 | 139605.506 | 77.6627934 | **<0.00001** | Aridlands |
| (Intercept) | -16.079953 | 0.23786404 | 59.3206275 | -67.601446 | **<0.00001** | Western Forest |
| year_since_1969 | 0.0249044 | 2.29E-04 | 214624.291 | 108.571773 | **<0.00001** | Western Forest |
| (Intercept) | -16.496221 | 0.42703138 | 29.535713 | -38.629998 | **<0.00001** | Boreal Forest |
| year_since_1969 | 0.02055381 | 2.60E-04 | 144129.764 | 78.9239838 | **<0.00001** | Boreal Forest |
| (Intercept) | -21.149892 | 0.19029974 | 88.2949882 | -111.13989 | **<0.00001** | Wetland |
| year_since_1969 | 0.07146027 | 1.90E-04 | 437624.317 | 376.781697 | **<0.00001** | Wetland |
| (Intercept) | -16.224505 | 0.60481837 | 22.0208031 | -26.825417 | **<0.00001** | Habitat Generalist |
| year_since_1969 | 0.03609136 | 1.77E-04 | 601827.075 | 203.382656 | **<0.00001** | Habitat Generalist |
| (Intercept) | -17.70426 | 0.57372769 | 7.88674417 | -30.858298 | **<0.00001** | Arctic Tundra |
| year_since_1969 | 0.03702689 | 0.00152971 | 1695.11876 | 24.2051941 | **<0.00001** | Arctic Tundra |
| (Intercept) | -16.625161 | 0.36579604 | 35.573756 | -45.449266 | **<0.00001** | Forest Generalist |
| year_since_1969 | 0.04268586 | 1.37E-04 | 686870.308 | 312.095903 | **<0.00001** | Forest Generalist |
| (Intercept) | -21.106998 | 1.04333342 | 3.77053432 | -20.230347 | **<0.00001** | Coasts |
| year_since_1969 | 0.09100524 | 0.00216601 | 1534.90013 | 42.0151681 | **<0.00001** | Coasts |
| (Intercept) | -15.408926 | 0.49326552 | 24.9097584 | -31.238603 | **<0.00001** | Grassland |
| year_since_1969 | 0.00477542 | 2.09E-04 | 412902.548 | 22.8267523 | 0.16057 | Grassland |
| (Intercept) | -17.350215 | 1.50982733 | 7.08618937 | -11.491523 | **<0.00001** | Introduced |
| year_since_1969 | 0.04985819 | 3.54E-04 | 219802.117 | 140.722753 | **<0.00001** | Introduced |

**Table S6.** Summary statistics for interaction potentials by predator species in Figure 2. Data included both recorded and predicted predator-prey interactions. Compare to Table 2, where functional compatibility does not scale with the number of species in a community.

| **Predator** | Mean of log-transformed temporal trends of interaction potentials | SD of log-transformed temporal trends of interaction potentials | Number of interaction pairs with decreasing temporal trends | Number of interaction pairs with increasing temporal trends |
| --- | --- | --- | --- | --- |
| *Falco sparverius* | -0.0102 | 0.0324 | 25 | 5 |
| *Buteo swainsoni* | 0.0142 | 0.0260 | 10 | 25 |
| *Buteo jamaicensis* | 0.0285 | 0.0304 | 10 | 68 |
| *Accipiter cooperii* | 0.0379 | 0.0316 | 9 | 163 |
| *Aquila chrysaetos* | 0.0166 | 0.0472 | 37 | 77 |
| *Falco mexicanus* | 0.0207 | 0.0210 | 8 | 50 |
| *Buteo regalis* | 0.0142 | 0.0253 | 6 | 21 |
| *Buteo lineatus* | 0.0488 | 0.0252 | 0 | 16 |
| *Haliaeetus leucocephalus* | 0.1089 | 0.0599 | 1 | 156 |
| *Buteo platypterus* | 0.0117 | 0.0322 | 14 | 21 |
| *Accipiter striatus* | 0.0174 | 0.0250 | 21 | 80 |
| *Accipiter atricapillus* | 0.0005 | 0.0311 | 35 | 40 |
| *Falco peregrinus* | 0.0663 | 0.0164 | 0 | 35 |
| *Falco columbarius* | 0.0517 | 0.0371 | 3 | 42 |

**Table S7.** Statistics of the linear model with spatial lag between turnover of predator-prey interactions and environmental changes between the years 1970 and 2021. These results are part of the sensitivity analysis, where we calculated interaction potentials using absolute instead of relative abundances. Turnover was calculated as the difference between matrices of interaction potentials, including both recorded and predicted predator-prey interactions. Environmental change includes changes in land use, mean annual temperature, and mean annual precipitation, and the interaction between changes in temperature and precipitation within each local community. See *Methods* for details on model fitting. Compare to Table 1’s results with relative abundances.

| **Environmental change** | **Estimate** | **SE** | ***z* value** | ***p* value** |
| --- | --- | --- | --- | --- |
| (Intercept) | 1.24448 | 0.29112 | 4.27484 | **0.00002** |
| Temperature | -0.01817 | 0.12301 | -0.14773 | 0.88256 |
| Precipitation | -0.00287 | 0.14317 | -0.02004 | 0.98401 |
| Settlements | 0.44601 | 0.16262 | 2.74262 | **0.00610** |
| Wild | -0.07161 | 0.26281 | -0.27246 | 0.78527 |
| Cultured | -0.01518 | 0.26501 | -0.05727 | 0.95433 |
| Temperature: precipitation | 0.12023 | 0.12800 | 0.93929 | 0.34758 |
| N = 1178, df = 1169; $\rho$ = 0.881, *p* value for $\rho$ < 0.0001; pseudo R^2^ = 0.423 | | | | |

**Table S8.** Statistics of linear mixed effects models for the temporal trends of predation risks for prey species by groups. These results are part of the sensitivity analysis, where we calculated interaction potentials using absolute instead of relative abundances. The upper table shows the results by functional groups, lower table shows the results by breeding biomes. Compare to Table S1’s results with relative abundances.

| **Parameters** | **Estimate** | **Std. Error** | **df** | **t value** | **Adjusted *p* value** | **Group** |
| --- | --- | --- | --- | --- | --- | --- |
| (Intercept) | -3.6927438 | 0.13090708 | 262.553382 | -28.208892 | **<0.00001** | landbird |
| year_since_1969 | 0.02252683 | 8.22E-05 | 2773345.25 | 273.970656 | **<0.00001** | landbird |
| (Intercept) | -7.3167315 | 0.37247959 | 12.529302 | -19.643309 | **<0.00001** | shorebird |
| year_since_1969 | 0.03010477 | 3.33E-04 | 92543.9874 | 90.3474414 | **<0.00001** | shorebird |
| (Intercept) | -9.2637264 | 0.24434754 | 43.8008072 | -37.912091 | **<0.00001** | waterbird |
| year_since_1969 | 0.06315476 | 3.29E-04 | 158603.227 | 192.124098 | **<0.00001** | waterbird |
| (Intercept) | -7.8335009 | 0.36630453 | 28.5603329 | -21.385214 | **<0.00001** | waterfowl |
| year_since_1969 | 0.06263147 | 2.43E-04 | 222495.701 | 257.781874 | **<0.00001** | waterfowl |
| (Intercept) | -5.0843109 | 1.64150944 | 6.99440884 | -3.0973388 | **0.01740462** | other |
| year_since_1969 | 0.02689909 | 3.18E-04 | 212110.091 | 84.6634812 | **<0.00001** | other |

| **parameters** | **Estimate** | **Std. Error** | **df** | **t value** | **Adjusted *p* value** | **Breeding biome** |
| --- | --- | --- | --- | --- | --- | --- |
| (Intercept) | -4.0815016 | 0.26601581 | 44.5779948 | -15.343079 | **<0.00001** | Eastern Forest |
| year_since_1969 | 0.0276104 | 1.54E-04 | 549791.063 | 178.773512 | **<0.00001** | Eastern Forest |
| (Intercept) | -4.8382436 | 0.37537216 | 41.0378614 | -12.889191 | **<0.00001** | Aridlands |
| year_since_1969 | 0.01574511 | 3.01E-04 | 141319.896 | 52.2553762 | **<0.00001** | Aridlands |
| (Intercept) | -4.3700749 | 0.22536225 | 61.1376164 | -19.391335 | **<0.00001** | Western Forest |
| year_since_1969 | 0.01752157 | 2.25E-04 | 220348.614 | 78.033093 | **<0.00001** | Western Forest |
| (Intercept) | -4.0581056 | 0.4285177 | 28.9344455 | -9.4701003 | **<0.00001** | Boreal Forest |
| year_since_1969 | 0.00964171 | 2.47E-04 | 160769.738 | 39.0657097 | **<0.00001** | Boreal Forest |
| (Intercept) | -8.7844245 | 0.20887007 | 82.4215614 | -42.056884 | **<0.00001** | Wetland |
| year_since_1969 | 0.06047161 | 1.86E-04 | 459345.644 | 325.142386 | **<0.00001** | Wetland |
| (Intercept) | -4.1094569 | 0.61022214 | 22.1908552 | -6.7343622 | **<0.00001** | Habitat Generalist |
| year_since_1969 | 0.02732142 | 1.76E-04 | 601150.4 | 155.491061 | **<0.00001** | Habitat Generalist |
| (Intercept) | -5.4684316 | 0.52551631 | 8.52651585 | -10.405827 | **<0.00001** | Arctic Tundra |
| year_since_1969 | 0.0295415 | 0.00143864 | 1695.10355 | 20.534344 | **<0.00001** | Arctic Tundra |
| (Intercept) | -4.5001501 | 0.36473431 | 35.8256537 | -12.338159 | **<0.00001** | Forest Generalist |
| year_since_1969 | 0.03404694 | 1.36E-04 | 691290.255 | 251.092501 | **<0.00001** | Forest Generalist |
| (Intercept) | -8.6647211 | 1.10381835 | 3.45716662 | -7.8497708 | **0.00254457** | Coasts |
| year_since_1969 | 0.06672674 | 0.00229363 | 1534.9571 | 29.0921444 | **<0.00001** | Coasts |
| (Intercept) | -3.2261123 | 0.4982699 | 24.6427172 | -6.474628 | **<0.00001** | Grassland |
| year_since_1969 | -0.0036441 | 2.07E-04 | 416279.93 | -17.563367 | **<0.00001** | Grassland |
| (Intercept) | -5.0843109 | 1.64150944 | 6.99440884 | -3.0973388 | **0.01740462** | Introduced |
| year_since_1969 | 0.02689909 | 3.18E-04 | 212110.091 | 84.6634812 | **<0.00001** | Introduced |

**Table S9.** Statistics for the latitudinal shifts of each predator, estimated from linear models between the latitudinal centroid and time (see *Methods*). These results are part of the sensitivity analysis, where we calculated interaction potentials using absolute instead of relative abundances. Compare to Table S2’s results with relative abundances.

| **Estimate** | **SE** | **statistic** | **Predator** | **Adjusted *p* value** |
| --- | --- | --- | --- | --- |
| -0.0105822 | 0.00279373 | -3.7878272 | *Falco sparverius* | **0.000626** |
| -0.0343366 | 0.0072638 | -4.7270902 | *Buteo swainsoni* | **0.0000327** |
| 0.00712487 | 0.00260517 | 2.73489548 | *Buteo jamaicensis* | **0.0113** |
| -0.0208016 | 0.00152664 | -13.625711 | *Accipiter cooperii* | **<0.00001** |
| 0.00887031 | 0.00113572 | 7.81028866 | *Aquila chrysaetos* | **<0.00001** |
| 0.006317 | 1.09E-04 | 57.7929878 | *Falco mexicanus* | **<0.00001** |
| 0.02109376 | 0.00346852 | 6.08148815 | *Buteo regalis* | **<0.00001** |
| 0.03764414 | 0.00173968 | 21.6385166 | *Buteo lineatus* | **<0.00001** |
| -0.1005975 | 0.00389444 | -25.831058 | *Haliaeetus leucocephalus* | **<0.00001** |
| -5.61E-04 | 0.00312569 | -0.1793258 | *Buteo platypterus* | 0.87631146 |
| 0.00537065 | 0.00143116 | 3.75264852 | *Accipiter striatus* | **<0.00001** |
| -0.0061741 | 2.75E-04 | -22.475159 | *Accipiter atricapillus* | **<0.00001** |
| 0.01006475 | 0.00316157 | 3.18346358 | *Falco peregrinus* | **0.00352** |
| -0.0071981 | 7.78E-04 | -9.2526394 | *Falco columbarius* | **<0.00001** |
